# The conserved endocannabinoid anandamide modulates olfactory sensitivity to induce hedonic feeding in *C. elegans*

**DOI:** 10.1101/2021.05.13.444082

**Authors:** Anastasia Levichev, Serge Faumont, Rachel Z. Berner, Zhifeng Purcell, Shawn R. Lockery

## Abstract

The ability of cannabis to increase consumption of food has been known for centuries. In addition to producing hyperphagia, cannabinoids can amplify existing preferences for calorically dense, palatable food sources, a phenomenon called hedonic feeding. These effects result from the action of plant-derived cannabinoids on brain receptors where they mimic natural ligands called endocannabinoids. The high degree of conservation of cannabinoid signaling at the molecular level across the animal kingdom suggests hedonic feeding may also be widely conserved. Here we show that exposure of *C. elegans* to anandamide, an endocannabinoid common to nematodes and mammals, shifts both appetitive and consummatory responses toward nutritionally superior food, an effect analogous to hedonic feeding. We find that anandamide’s effect on feeding requires the *C. elegans* cannabinoid receptor NPR-19 but it can also be mediated by the human CB1 cannabinoid receptor, indicating functional conservation between the nematode and mammalian endocannabinoid systems for regulation of food preferences. Furthermore, the effect of anandamide in *C. elegans* is bidirectional, as it increases appetitive and consummatory responses to superior food but decreases these responses to inferior food. This bidirectionality is mirrored at the cellular level. Anandamide’s behavioral effects require the AWC chemosensory neurons, and anandamide renders these neurons more sensitive to superior food and less sensitive to inferior food. Our findings reveal a surprising degree of functional conservation in the effects of endocannabinoids on hedonic feeding across species and establish a new system in which to investigate the cellular and molecular basis of endocannabinoid system function in the regulation of food choice.

## Introduction

It has been known for centuries that smoking or ingesting preparations of the plant *Cannabis sativa* stimulates appetite (Abel, 1971; Kirkham & Williams, 2001). Users report persistent hunger while intoxicated, even if previously satiated. This feeling of hunger is often accompanied by a strong and specific desire for foods that are sweet or high in fat content, a phenomenon colloquially known as “the munchies” (Abel, 1975; Foltin et al., 1986, 1988; Halikas et al., 1971; Hollister, 1971; Tart, 1970). The effects of cannabinoids on appetite result mainly from Δ^9^-tetrahydrobannabinol (THC), a plant-derived cannabinoid. THC acts at cannabinoid receptors in the brain where it mimics endogenous ligands called endocannabinoids, which include N-arachidonoylethanolamine (AEA) and 2-arachidonoylglycerol (2-AG). AEA and 2-AG are the best studied signaling molecules of the mammalian endocannabinoid system, which comprises the cannabinoid receptors CB1 and CB2, metabolic enzymes for synthesis and degradation of the endocannabinoids, and a variety of ancillary proteins involved in receptor trafficking and modulation (Bauer et al., 2012; Fu et al., 2011; Jin et al., 1999; Kaczocha et al., 2009, 2012; Liedhegner et al., 2014; Martini et al., 2007; Oddi et al., 2009; Rozenfeld & Devi, 2008).

A large number of studies in laboratory animals have established a strong link between endocannabinoid signaling and energy homeostasis, defined as the precise matching of caloric intake with energy expenditure to maintain body weight (Cristino et al., 2014). Food deprivation increases endocannabinoid levels in the limbic forebrain, which includes the nucleus accumbens and hypothalamus, two brain regions that express CB1 receptors and contribute to the appetitive drive for food (Kirkham et al., 2002). Systemic administration of THC or endogenous cannabinoids increases feeding (Williams & Kirkham, 1999). Similarly, micro-injection of cannabinoid receptor agonists or endocannabinoids directly into the nucleus accumbens also increases feeding (Deshmukh & Sharma, 2012; Mahler et al., 2007). Thus, the endocannabinoid system can be viewed as a short-latency effector system for restoring energy homeostasis under conditions of food deprivation (Cristino et al., 2014; Devane et al., 1988; Munro et al., 1993; Parker, 2017).

To respond effectively to an energy deficit, an animal should be driven both to seek food (*appetitive* behavior) and, once food is encountered, to maximize caloric intake (*consummatory* behavior). The endocannabinoid system is capable of orchestrating both aspects of this response simultaneously. With respect to appetitive behavior, CB1 agonists reduce the latency to feed (Freedland et al., 2000; Gallate et al., 1999; Gallate & McGregor, 1999; Maccioni et al., 2008; McLaughlin et al., 2003; Salamone et al., 2007; Thornton-Jones et al., 2005) and induce animals to expend more effort to obtain a given food or liquid reward (Barbano et al., 2009; Freedland et al., 2000; Gallate et al., 1999; Guegan et al., 2013), whereas CB1 antagonists have the opposite effect (Freedland et al., 2000; Gallate et al., 1999; Gallate & McGregor, 1999; Maccioni et al., 2008; McLaughlin et al., 2003; Salamone et al., 2007; Thornton-Jones et al., 2005). With respect to consummatory behavior, studies in rodents show that administration of THC or endocannabinoids specifically alters food preferences in favor of palatable, calorically dense foods, such as those laden with sugars and fats, as opposed to laboratory pellets. For example, THC causes rats to consume larger quantities of chocolate cake batter without affecting consumption of simultaneously available laboratory pellets (Koch & Matthews, 2001). It also causes them to consume larger quantities of sugar water than plain water, and of dry pellets than watered-down pellet mash, which is calorically dilute (Brown et al., 1977). Administration of endocannabinoids, including microinjection into the nucleus accumbens, has similar effects, which can be blocked by simultaneous administration of CB1 antagonists (Deshmukh & Sharma, 2012; Escartín-Pérez et al., 2009a; Shinohara et al., 2009). CB1 antagonists, administered alone, specifically suppress consumption of sweet and fatty foods in rats (Arnone et al., 1997; Gessa et al., 2006; Mathes et al., 2008) as well as in primates (Simiand et al., 1998), indicating that basal endocannabinoid titers can be regulated up or down to re-establish energy homeostasis.

There is considerable support for the hypothesis that animals treated with cannabinoids consume larger quantities of calorically dense foods because cannabinoids amplify the pleasurable or rewarding aspects of these foods. This phenomenon has been termed *hedonic amplification* (Castro & Berridge, 2017; Mahler et al., 2007), whereas the food-specific increase in consumption it engenders has been termed *hedonic feeding* (Edwards & Abizaid, 2016). Inferences concerning pleasurable and rewarding aspects of animal experience can be difficult to establish, but both THC and AEA specifically increase the vigor of licking at spouts delivering sweet fluids (Davis & Smith, 1992; Higgs et al., 2003). In a more direct measure of hedonic responses, the frequency of orofacial movements previously shown to be associated with highly preferred foods can be monitored in response to oral delivery of a sucrose solution (Grill & Norgren, 1978). Injection of THC or a CB1 antagonist respectively increases or decreases this frequency (Jarrett et al., 2005), suggesting that pleasure may have been increased by cannabinoid administration.

Cannabinoid effects on hedonic responses may be at least partially chemosensory in origin, including both taste (gustation) and smell (olfaction). With respect to gustation, a majority of sweet-sensitive taste cells in the mouse tongue are immunoreactive to CB1, and a similar proportion shows increased response to saccharin, sucrose, and glucose following endocannabinoid administration (Yoshida et al., 2010, 2013). These effects are recapitulated in afferent nerves from the tongue (Yoshida et al., 2010), as administration of AEA or 2-AG specifically increases chorda tympani responses to sweeteners rather than NaCl (salt), HCl (sour), quinine (bitter), or monosodium glutamate (umami). With respect to olfaction, CB1 receptors expressed in the olfactory bulb are required for post-fasting hyperphagia in mice, and THC decreases the threshold of food-odor detection during exploratory behavior (Soria-Gómez et al., 2014).

The high degree of conservation of the endocannabinoid system at the molecular level is well established (Elphick, 2012). Although CB1 and CB2 receptors are unique to chordates, there are numerous candidates for cannabinoid receptors in most animals. Furthermore, orthologs of the enzymes involved in biosynthesis and degradation of endocannabinoids occur throughout the animal kingdom. This degree of molecular conservation, coupled with the universal need in all organisms to regulate energy balance, suggests the hypothesis that hedonic amplification and hedonic feeding are also widely conserved, but studies in animals other than rodents and primates appear to be lacking.

The present study tests the hypothesis that the hedonic effects of cannabinoids are conserved in the nematode *C. elegans*. This organism diverged from the line leading to mammals more than 500 million years ago (Raible & Arendt, 2004). Nevertheless, *C. elegans* has a fully elaborated endocannabinoid signaling system including: (i) a functionally validated endocannabinoid receptor NPR-19, which is encoded by the gene *npr-19* (Oakes et al., 2017); (ii) the endocannabinoids AEA and 2-AG, which it shares with mammals (Higgs et al., 2003; Lehtonen et al., 2008, 2011; Sugiura et al., 1995), (iii) orthologs of the mammalian endocannabinoid synthesis enzymes NAPE-1 and NAPE-2, and DAGL (Harrison et al., 2014), and (iv) orthologs of endocannabinoid degradative enzymes FAAH and MAGL (Y97E10AL.2 in worms) (Oakes et al., 2017). Endocannabinoid signaling in *C. elegans* is so far known to contribute to six main phenotypes: (i) axon navigation during regeneration (Pastuhov et al., 2012, 2016), (ii) lifespan regulation related to dietary restriction (Harrison et al., 2014; Lucanic et al., 2011) (iii) altered progression through developmental stages (Harrison et al., 2014; Reis-Rodrigues et al., 2016), (iv) suppression of nociceptive withdrawal responses (Oakes et al., 2017), (v) inhibition of feeding rate (Oakes et al., 2017), and (vi) inhibition of locomotion (Oakes et al., 2017, 2019). Despite considerable conservation between the *C. elegans* and mammalian endocannabinoid systems, to our knowledge the effects of cannabinoids on food preference in *C. elegans* have not been described.

The feeding ecology of *C. elegans* supports the possibility of hedonic feeding in this organism. *C. elegans* feeds on bacteria in decaying plant matter (Frézal & Félix, 2015). It finds bacteria by chemotaxis driven by a combination of gustatory and olfactory cues (Bargmann et al., 1993; Bargmann & Horvitz, 1991). Bacteria are ingested through the worm’s pharynx, a rhythmically active muscular pump that constitutes the animal’s throat. Although *C. elegans* is an omnivorous bacterivore, different species of bacteria have a characteristic quality as a food source defined by the rate of growth of individual worms feeding on that species (Δ length/unit time). Hatchlings are naïve to food quality but in a matter of hours begin to exhibit a preference for nutritionally superior species (*favored*) over nutritionally inferior species (*non-favored*) (Shtonda, 2006).

Here we show that transient exposure of *C. elegans* to the endocannabinoid AEA simultaneously biases appetitive and consummatory responses toward favored food. With respect to appetitive responses, the fraction of worms approaching and dwelling on patches of favored food increases whereas the fraction approaching and dwelling on non-favored food decreases. With respect to consummatory responses, feeding rate in favored food increases whereas feeding rate in non-favored food decreases. Taken together, the appetite and consummatory manifestations of cannabinoid exposure in *C. elegans* imply increased consumption of favored food characteristic of hedonic feeding. We also find that AEA’s effects require the NPR-19 cannabinoid receptor. Further, AEA’s effects persist when *npr-19* is replaced by the human CB1 receptor gene CNR1, indicating a high degree of conservation between the nematode and mammalian endocannabinoid systems. At the neuronal level, we find that under the influence of AEA, AWC, a primary olfactory neuron required for chemotaxis to food, becomes more sensitive to favored food and less sensitive to non-favored food. Together, our findings indicate that the hedonic effects of endocannabinoids are conserved in *C. elegans*.

## Results

### AEA exposure increases preference for favored food

We pre-exposed well-fed, adult, wild type (N2 Bristol) worms to the endocannabinoid AEA by incubating them for 20 min at a concentration of 100 μM. Food preference was measured by placing a small population of worms at the starting point of a T-maze baited with patches of favored and non-favored bacteria at equal optical densities (OD_600_ 1), where optical density served as a proxy for bacteria concentration (see Materials and Methods; Fig. 1A). This assay is analogous to assays used in mammalian studies in which both palatable and standard food options are simultaneously available (Brown et al., 1977; Deshmukh & Sharma, 2012; Escartín-Pérez et al., 2009a; Koch & Matthews, 2001; Shinohara et al., 2009). The number of worms in each food patch was counted at 15-minute intervals for one hour. At each time point, we quantified preference in terms of the index *I* = (*n*_F_ − *n*_NF_)/(*n*_F_ + *n*_NF_), where *n*_F_ and *n*_NF_ are the number of worms in favored and non-favored food, respectively, and *I* = 0 indicates indifference between the two food types. We found that AEA exposure increased preference for favored food (Fig. 1B, C; Suppl. Table 1, line 2). This effect lasted at least 60 minutes without significant decrement (Fig. 1B; Suppl. Table 1, line 3-4) despite the absence of AEA on the assay plates. Thus, the amount of AEA absorbed by worms during the exposure period was sufficient to maintain the increased preference for favored food throughout the observation period.

**Fig 1.**
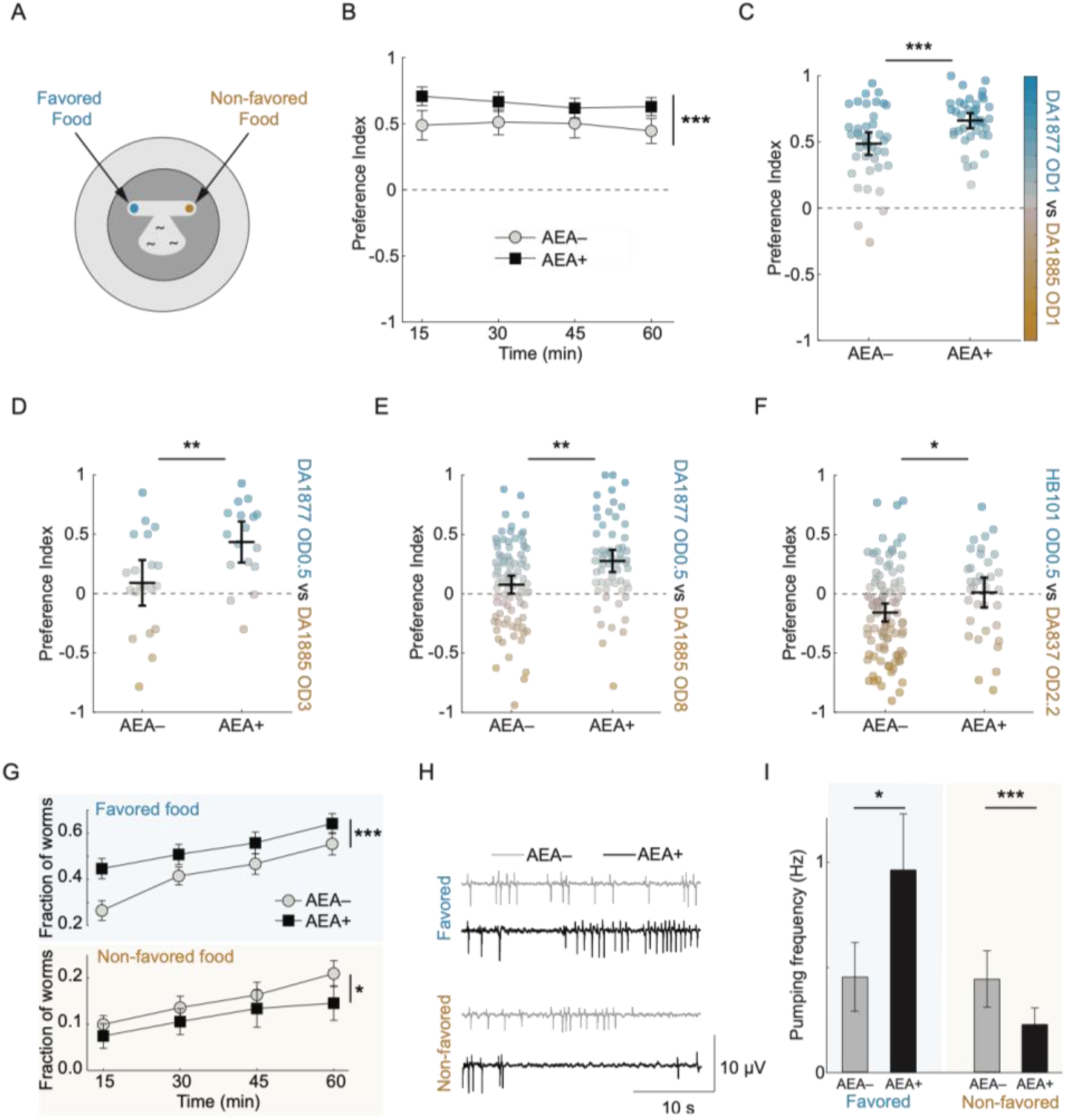
AEA-mediated hedonic feeding. **A**. Food preference assay. T-maze arms were baited with patches of favored (blue) and non-favored (orange) bacteria. **B**. Mean preference index (*I*) versus time for AEA-exposed animals (AEA+) and unexposed controls (AEA–), where *I* > 0 is preference for favored food, *I* < 0 is preference for non-favored food, and *I* = 0 is indifference (*dashed line*). Favored food, DA1877, OD 1; non-favored food, DA1885, OD 1. **C**. Summary of the data in **B**. Each dot is mean preference over time in a single T-maze assay. *Dot color* indicates preference index according to the color scale on the right. **D, E**. Effect of AEA on preference when baseline preference is at the indifference point (symbols as in **C**). For preference time courses, see Supp. Fig. 1. In **D**: Favored food, DA1877, OD 0.5; non-favored food, DA1885, OD 3. In **E**: favored food, DA1877, OD 0.5; non-favored food, DA1885, OD 8. **F**. Effect of AEA on preference for a different pair of favored and non-favored bacteria (symbols as in **C**). Favored food, HB101, OD 0.5; non-favored food, DA837, OD 2.2. For preference time course, see Supp. Fig. 1. **G**. Effect of AEA on fraction of worms in favored and non-favored food patches versus time. Same experiment as in panels **B, C. H, I**. Effect of AEA on pharyngeal pumping in favored versus non-favored food. Favored food, DA1877, OD 0.8; non-favored food, DA1885, OD 0.8. **H** shows electrical recordings of four individual worms under the conditions shown. Each spike is the electrical correlate of one pump. Traces were selected to represent the population median pumping frequency in each condition. **I** shows mean pumping frequency in each condition. For statistics in **B**-**G** and **I**, see Supp. Table 1. Symbols: *, *p* < 0.05; **, *p* < 0.01; ***, *p* < 0.001; n.s., not significant. Error bars, 95% confidence interval.

A simple interpretation of the data in Fig. 1B, C is that AEA exposure specifically increases the relative attractiveness of favored food. However, an alternative interpretation is that AEA promotes the attractiveness of whichever food is already preferred under the baseline conditions of the experiment (AEA–). To test this possibility, we titrated the densities of favored and non-favored food so that under baseline conditions neither food was preferred (*I* ≈ 0; Fig. 1D, E; Suppl. Fig. 1A, B). Under these conditions, AEA still increased the preference for favored food (Suppl. Table 1, line 6, 10). This finding supports the hypothesis that AEA differentially affects accumulation based on food identity, not relative food density. Finally, we found that AEA’s effect on preference generalized to a different pair of favored and non-favored bacteria (Fig. 1F; Suppl. Fig. 1C; Suppl. Table 1, line 14). Taken together, the data in Fig. 1B-F show that AEA’s ability to increase preference for favored food is not limited to a particular pair of foods or their relative concentrations.

In mammals, cannabinoid administration can differentially increase responses to favored versus non-favored food. Because worms in the T-maze assay could occupy foodless regions of the assay plate in addition to the food patches themselves, the increased accumulation in favored food could represent an increased appetitive response to favored food, a decreased appetitive response to non-favored food, or both. Further analysis revealed that AEA exposure increased the fraction of worms in favored food and decreased the fraction in non-favored food (Fig. 1G; Suppl. Table 1, line 18, 22). Thus, AEA exposure produces a bidirectional effect on appetitive responses to favored versus non-favored food, the net result of which is increased accumulation in favored food.

Are these food-specific appetitive responses accompanied by food-specific changes in consumption behavior? *C. elegans* swallows bacteria by means of rhythmic contractions of its pharynx, a muscular organ comprising its throat; each contraction is called a pump. We recorded pumping electrically in individual worms restrained in a microfluidic channel with integrated electrodes (Lockery et al., 2012; David M. Raizen & Avery, 1994). The channel contained either favored or non-favored food and pumping was recorded for 1 min following a 3 min accommodation period. Under these conditions, pumping rate is a reasonable proxy for the amount of food consumed because food concentration at this optical density is effectively constant. Unexposed worms pumped at equal frequencies in the presence of favored and non-favored species (Fig. 1H, I; Suppl. Table 1, line 25). However, under the influence of AEA, pumping frequency in favored food increased whereas pumping frequency in non-favored food decreased (Fig. 1H, I; Suppl. Table 1, line 26-27). Thus, the effects of AEA exposure on food consumption mirror its bidirectional effects on accumulation shown in Fig. 1G.

Taken together, the results in Fig. 1 demonstrate clear homologies between the effects of cannabinoids on feeding behavior in nematodes and mammals in two key respects. First, AEA differentially alters *appetitive* responses to favored and non-favored food, causing more worms to accumulate in the former and fewer in the latter. Second, AEA differentially alters *consummatory* responses measured in terms of feeding rate, causing individual worms to consume more favored food and less non-favored food per unit time. The appetitive and consummatory effects of AEA, acting in concert, are consistent with a selective increase in consumption of favored food, which is phenomenologically analogous to hedonic feeding in mammals (Edwards & Abizaid, 2016).

### AEA differentially modulates chemosensory responses to favored and non-favored food

In theoretical terms, accumulation in a food patch is determined by just two factors: entry rate and exit rate. Previous studies in *C. elegans* have shown that both rates can contribute to differential accumulation in one food versus another (Shtonda, 2006). Thus, AEA could modulate appetitive responses by acting on entry, exit rate, or both. Chemotaxis toward food patches is driven by olfactory neurons responding to airborne cues encountered at a distance (Bargmann et al., 1993; Bargmann & Horvitz, 1991). Thus, changes in entry rate might implicate changes in the function of olfactory neurons. A simple but powerful way to examine the contribution of entry rate is to spike food patches with a paralytic agent so worms that enter a patch cannot leave, thereby setting exit rate to zero. Under these conditions, if AEA exposure still modulates relative preference for favored versus non-favored food, then AEA must be differentially altering the entry rate into the two foods. To test this, we added sodium azide, a paralytic agent commonly used to immobilize nematodes (Hart, 2006), to both food patches in the T-maze. We found that AEA still produced a marked increase in preference for favored food (Fig. 2A; Suppl. Table 2, line 2), showing that it differentially affects patch entry rates.

**Fig 2.**
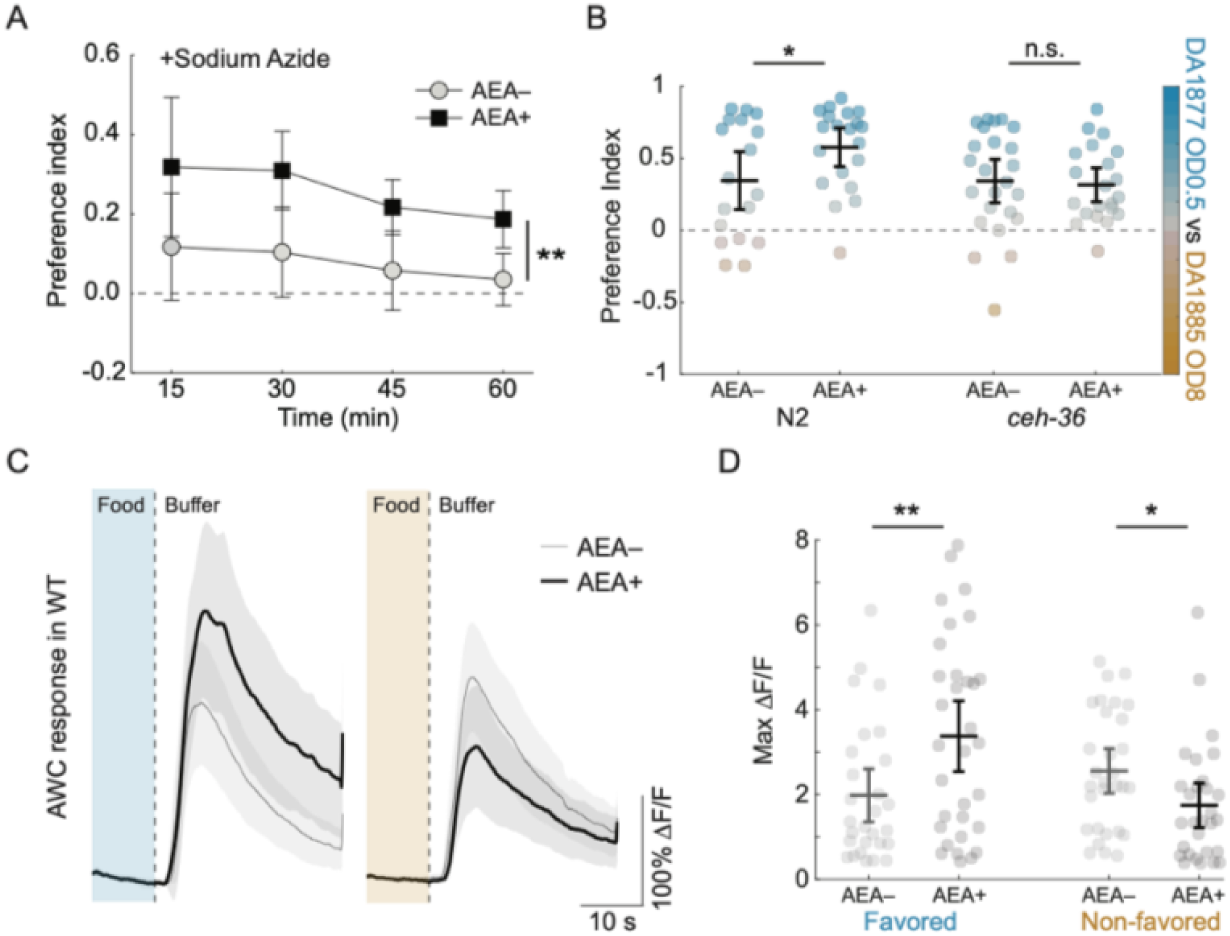
Chemosensory correlate of hedonic feeding. **A**. Mean preference index (*I*) versus time for AEA-exposed animals (AEA+) and unexposed controls (AEA–) when sodium azide was added to food patches. Favored food, DA1877, OD 0.5; non-favored food, DA1885, OD 3. **B**. Effect of AEA on preference in wild type (N2) and *ceh-36* mutants. Favored food, DA1877, OD 0.5; non-favored DA1885, OD 8. Each dot is mean preference in a single T-maze assay. **C**. Effect of AEA on the response of AWC neurons to the removal of favored or non-favored food. Each trace is average normalized fluorescence change (Δ*F*/*F*) versus time. Favored food (blue), DA1877, OD 1; non-favored food (orange), DA1885, OD 1. **D**. Summary of the data in in C, showing mean peak Δ*F*/*F*. For statistics in **A**-**D**, see Supp. Table 2. Symbols: *, *p* < 0.05; **, *p* < 0.01; n.s., not significant. Error bars and shading, 95% confidence interval.

Having found that AEA alters food-patch entry rates, we next considered the possibility that AEA acts on olfactory neurons to produce the appetitive component of hedonic feeding. *C. elegans* senses food or food-related compounds by means of 11 classes of chemosensory neurons (two neurons/class), which have sensory endings in the anterior sensilla near the mouth (Bargmann et al., 1993; Zaslaver et al., 2015). We focused on the AWC class, a pair of olfactory neurons that responds directly to many volatile odors (Leinwand et al., 2015) and is required for chemotaxis to them (Bargmann et al., 1993). To investigate whether AEA acts on AWC to alter food preference, we measured AEA’s effect on preference in *ceh-36* mutants, in which AWC function is selectively impaired. This gene is expressed only in AWC and the gustatory neuron class ASE. *ceh-36* is required for normal expression levels of genes essential for chemosensory transduction, particularly in AWC (Koga & Ohshima, 2004; Lanjuin et al., 2003). Accordingly, *ceh-36* mutants are strongly defective in their chemotaxis responses to three food-related odorants that directly activate AWC (Lanjuin et al., 2003). Although ASE neurons are required for chemotaxis to at least one AWC-sensed odorant (Leinwand et al., 2015), they do not respond directly to these compounds; rather, they inherit their response via peptidergic signaling from AWC. Thus, loss of appetitive responses in *ceh-36* mutants can be attributed to AWC neurons.

In T-maze assays, we found a modest strain × AEA interaction (*p* = 0.08), and a significant effect of AEA in wild type animals which was absent in the mutants (Fig. 2B; Suppl. Fig. 2A, B; Suppl. Table 2, line 6, 10-11, 13). This finding indicates that AWC is required for the appetitive component of hedonic feeding. With respect to the consummatory component, whereas AEA exposure had no effect on pumping frequency of *ceh-36* null worms in non-favored food, it still increased pumping frequency in favored food, just as it did in wild type worms (Suppl. Fig. 3, Suppl. Table 5, line 1-2), indicating that *ceh-36* is partially required for the consummatory component of hedonic feeding. Taken together, these data suggest that AWC is required for the normal magnitude of both components of hedonic feeding.

AWC is activated by *decreases* in the concentration of food or food-related odors (Calhoun et al., 2015; Chalasani et al., 2007; Zaslaver et al., 2015). AWC can nevertheless promote *attraction* to food patches because its activation truncates locomotory head bends away from the odor source, thereby steering the animal toward the odor source. Additionally, its activation causes the animal to stop moving forward, reverse, and resume locomotion in a new direction better aligned with the source; this behavioral motif is known as a pirouette (Pierce-Shimomura et al., 1999). To test whether AEA alters AWC sensitivity to favored and non-favored food, we compared AWC calcium transients in response to the removal of either type of food in wild type worms exposed to AEA, and in unexposed controls. In unexposed animals, AWC neurons responded equally to the removal of either food (Fig. 2C, D, Suppl. Table 2, line 21). However, exposure to AEA caused a dramatic change in food sensitivity, increasing AWC’s response to the removal of favored food and decreasing its response to the removal of non-favored food (Fig. 2C, D, Suppl. Table 2, line 17, 19-20, 22). This bidirectional effect mirrors AEA’s effect on both the appetitive and consummatory aspects of hedonic feeding (Fig. 1G, I) and is consistent with a model in which hedonic feeding is triggered at least in part by modulation of chemosensation in AWC neurons.

### Dissection of signaling pathways required for hedonic feeding

The NPR-19 receptor has been shown to be required for AEA-mediated suppression of withdrawal responses and feeding rate (Oakes et al., 2017). To test whether *npr-19* is required for hedonic feeding, we measured food preference in *npr-19* null mutants following exposure to AEA. Mutant worms failed to exhibit increased preference for favored food (Fig. 3A; Suppl. Fig. 2C, D; Suppl. Table 3, line 6-7). This defect was rescued by over-expressing *npr-19* under control of the native *npr-19* promoter (Fig. 3A; Suppl. Fig. 2C, E; Suppl. Table 3, line 11-12, 15-16, 18). We conclude that *npr-19* is required for the appetitive component of hedonic feeding. This defect was also rescued by over-expressing the human cannabinoid receptor CB1 under the same promoter (Fig. 3A; Suppl. Fig. 2F; Suppl. Table 3, line 20-21, 24-25, 27). This finding indicates a remarkable degree of conservation between the nematode and human endocannabinoid systems. With respect to the consummatory component of hedonic feeding, the role of *npr-19* was unclear: *npr-19* mutants worms exhibited only a partial phenotype which was not rescued by overexpression of either *npr-19* or CNR1 (Suppl. Fig. 3), despite evidence of rescue in a previous study (Oakes et al., 2017). Significant differences in experimental approach might explain this discrepancy (see Materials and Methods).

**Fig 3.**
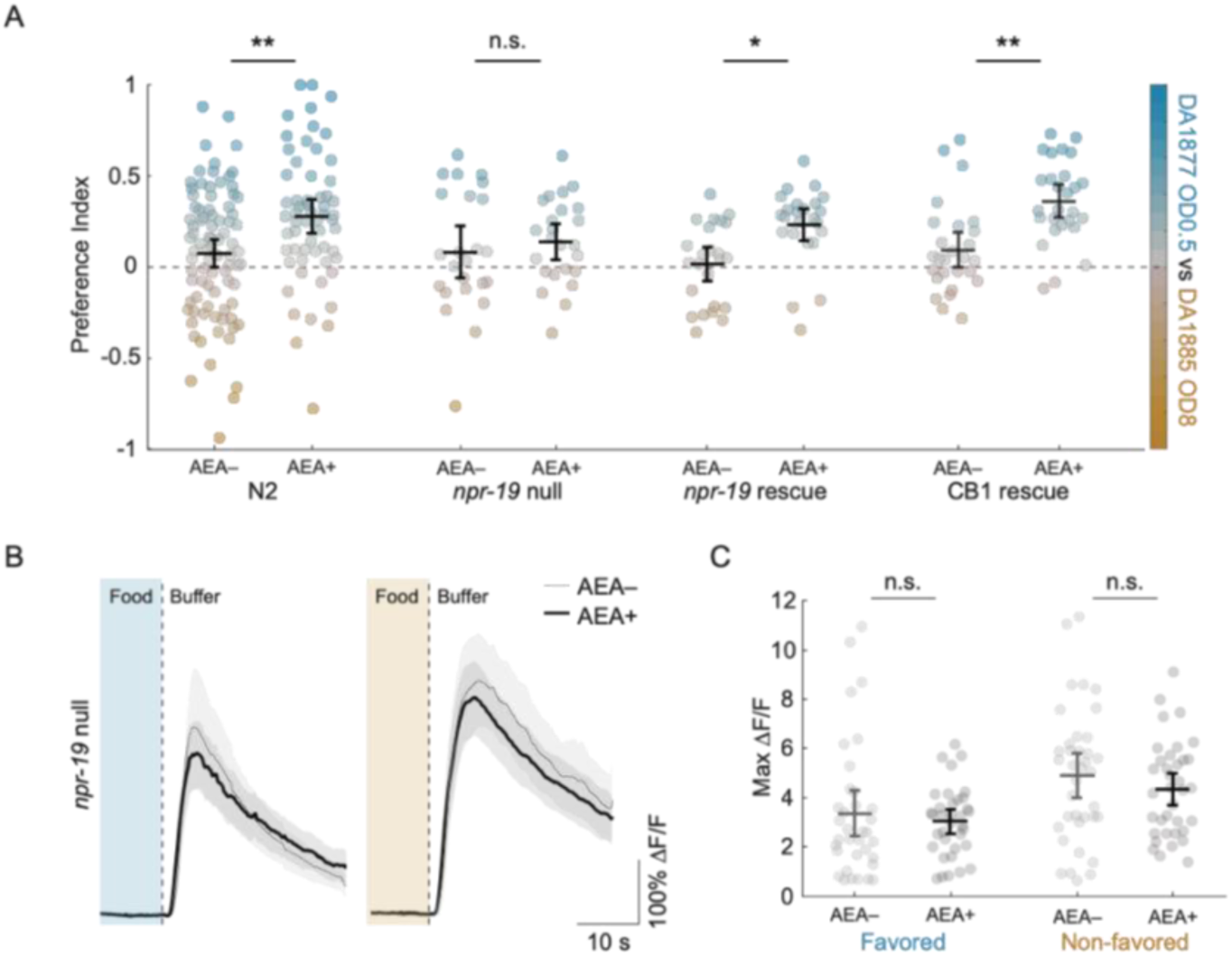
Requirement of NPR-19 for hedonic feeding and chemosensory modulation. **A**. Effect of AEA on preference in wild type worms (N2) and the indicated genetic background. Favored food, DA1877, OD 0.5; non-favored food, DA1885, OD 8. Each dot is mean preference over time in a single T-maze assay. *Dot color* indicates preference index according to the color scale on the right. **B**. Effect of AEA on the response of AWC neurons to the removal of favored or non-favored food in *npr-19* mutants. Each trace is average normalized fluorescence change (Δ*F*/*F*) versus time. Favored food (blue), DA1877, OD 1; non-favored food (orange), DA1885, OD 1. **C**. Summary of the data in **B**, showing mean peak Δ*F*/*F*. For statistics in **A**-**C**, see Supp. Table 3. Symbols: *, *p* < 0.05; **, *p* < 0.01; n.s., not significant. Error bars and shading, 95% confidence interval.

The forgoing results suggest a model of hedonic feeding in *C. elegans* in which activation of the NPR-19 receptor by AEA triggers a bidirectional change in AWC’s food sensitivity (Fig. 2C, D) to induce the appetitive component of hedonic feeding. We therefore tested whether *npr-19* is required for AEA’s effects on AWC. The effect of AEA on AWC’s response to food was abolished in *npr-19* mutants (Fig. 3B, C, Suppl. Table 3, line 30, 33-34, 39, 42-43). This phenotype was partially rescued by over-expression of the CB1 receptor (Suppl. Fig. 4A, B, Suppl. Table 5, line 12, 15, 18, 22, 24). We conclude that the appetitive component of AEA-induced hedonic feeding requires both the NPR-19 receptor and AWC neurons.

In perhaps the simplest model of AEA’s effect on AWC, NPR-19 is expressed in AWC, and activation of NPR-19 produces the observed bidirectional modulation of sensitivity to favored and non-favored food. To test this model, we characterized the *npr-19* expression pattern. This was done by expressing a p*npr-19*::GFP transgene together with either p*cho-1*::mCherry or p*eat-4*::mCherry, two neuronal markers whose expression pattern has been thoroughly characterized (Pereira et al., 2015; Serrano-Saiz et al., 2013). We observed expression of *npr-19* in body wall muscles together with an average of 29 neuronal somata in the head and 8 in the tail (Fig. 4A, Suppl. Table 6). Using positional cues in addition to the markers, we identified 28 of the GFP-positive somata, which fell into 15 neuron classes (Table 1). These classes could be organized into four functional groups: sensory neurons (URX, ASG, AWA, and PHC), interneurons (RIA, RIM, and LUA), motor neurons (URA and PDA), and pharyngeal neurons (M1, M3, MI, MC, I2, and I4). Although AWC could be identified in every worm by its characteristic position in the p*eat-4*::mCherry expressing strain, GFP expression was never observed in this neuron class. Our expression data, together with the absence of significant *npr-19* expression in AWC in RNA sequencing experiments based on the *C. elegans* Neuronal Gene Expression Map & Network (CeNGEN) consortium (Hammarlund et al., 2018), suggests that AWC does not express *npr-19*. These findings are inconsistent with a direct action of AEA on AWC neurons mediated by the NPR-19 receptor.

**Fig 4.**
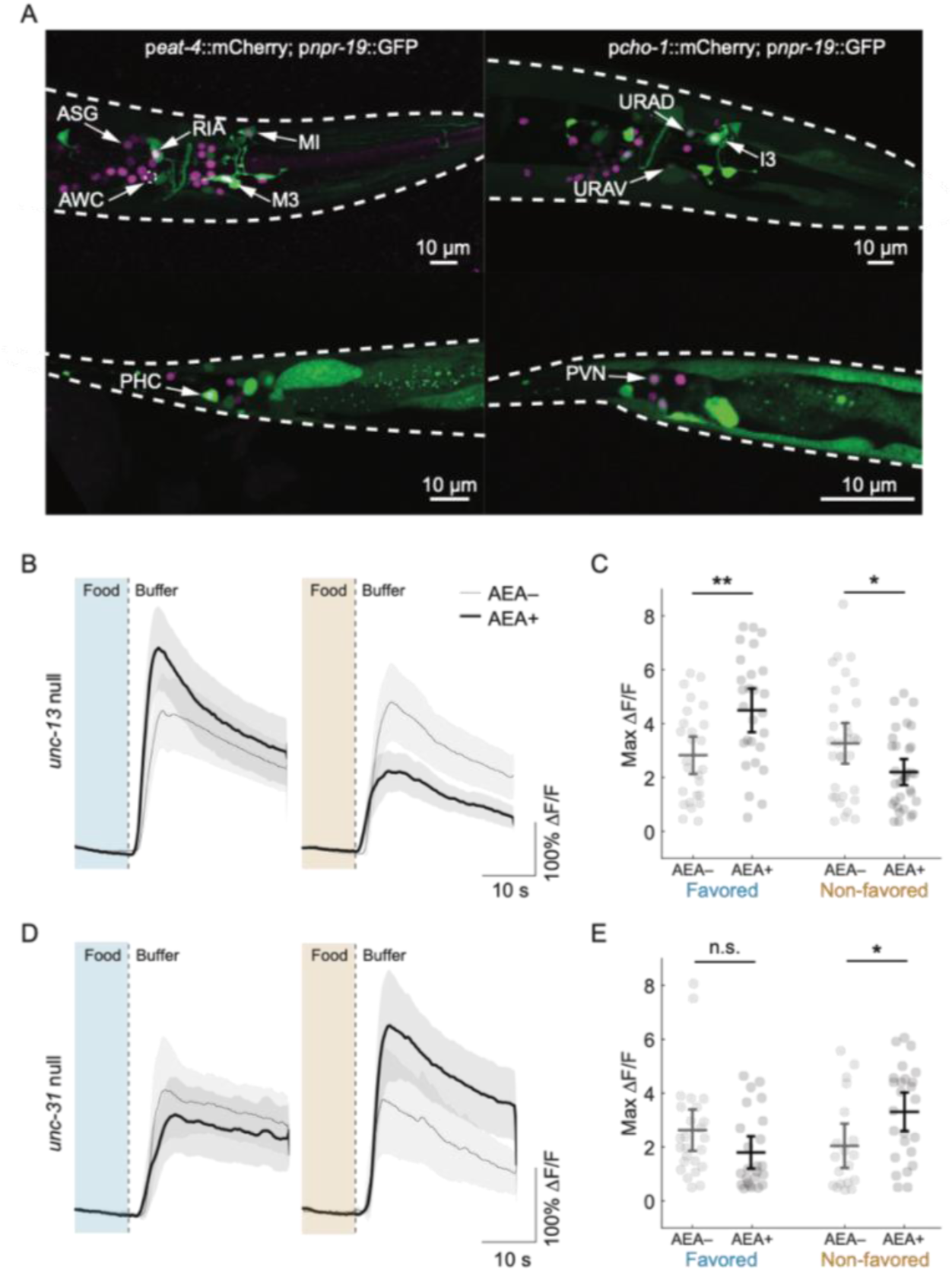
Genetic pathways underlying AEA-mediated AWC modulation. **A**. Expression pattern of *npr-19*. Green cells express *npr-19. Left*, magenta indicates expression of *eat-4*, a marker for glutamatergic neurons. *Dashed circle*, the soma of AWC, which is glutamatergic. *Right*, magenta indicates expression of *cho-1*, a marker for cholinergic neurons. *Top, bottom*, head and tail expression, respectively. **B**. Effect of AEA on the response of AWC neurons to the removal of favored or non-favored food in *unc-13* mutants. Each trace is average normalized fluorescence change (Δ*F*/*F*) versus time. Favored food (blue), DA1877, OD 1; non-favored food (orange), DA1885, OD 1. **C**. Summary of the data in **B**, showing mean peak Δ*F*/*F*. **D**. Effect of AEA on the response of AWC neurons to the removal of favored or non-favored food in *unc-31* mutants. Each trace is average normalized fluorescence change (Δ*F*/*F*) versus time. Favored food (blue), DA1877, OD 1; non-favored food (orange), DA1885, OD 1. **E**. Summary of the data in **D**, showing mean peak Δ*F*/*F*. For statistics in **B**-**E**, see Supp. Table 4. Symbols: *, *p* < 0.05; **, *p* < 0.01; n.s., not significant. Error bars and shading, 95% confidence interval.

**Table 1.**
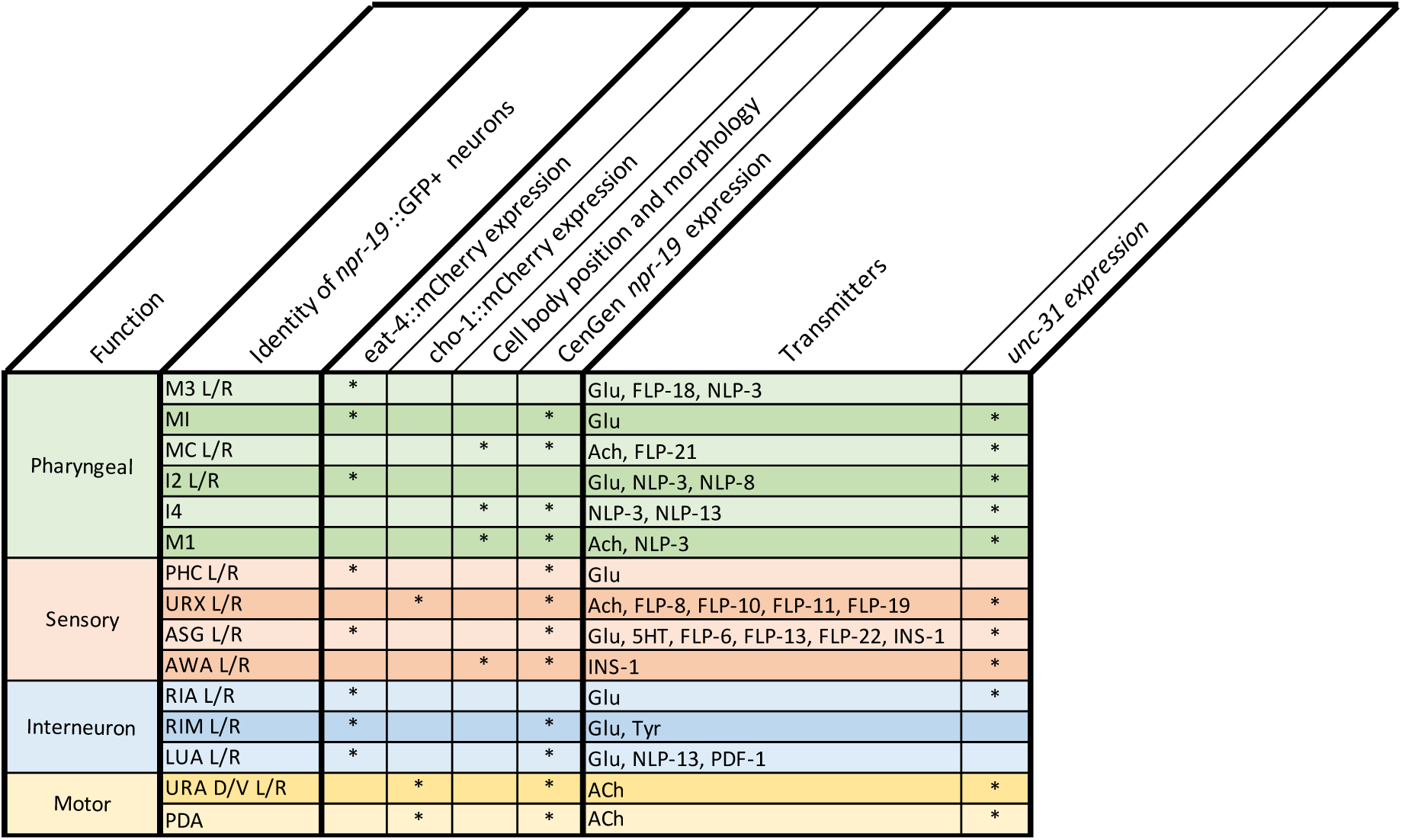
*npr-19*-expressing neurons. The *npr-19* expression pattern was characterized by expressing a p*npr-19*::GFP transgene together with either p*cho-1*::mCherry or p*eat-4*::mCherry, respectively labeling previously identified cholinergic and glutamatergic neurons (Pereira et al., 2015; Serrano-Saiz et al., 2013). GFP-positive neurons that expressed neither of the markers were identified by position and morphology, and confirmed by cross-reference to CeNGEN expression data showing *npr-19*. Also shown are neurotransmitter identity (Loer and Rand, 2016; Altun, 2011) and *unc-31* expression (CeNGEN) of each identified neuron class. See also Supp. Table 6.

The *npr-19* expression pattern supports at least two indirect models of AEA’s effect on AWC. In the first model, AWC inherits its sensitivity to AEA from incoming, AEA-sensitive, classical synaptic pathways (i.e., those that do not involve neuromodulatory transmitters). For example, in one common endocannabionoid signaling motif, endocannabinoids act as retrograde signals released by a postsynaptic neuron to suppress transmitter release by binding to cannabinoid receptors on presynaptic terminals. This motif could render AWC-related synaptic pathways sensitive to AEA. To determine whether this motif may be present in *C. elegans*, we searched the *C. elegans* connectome for the anatomical substrate of retrograde signaling: synaptically coupled pairs of neurons in which the *presynaptic* neuron expressed *npr-19* and the *postsynaptic* neuron expressed a key synthesis enzyme for AEA. The set of presynaptic, *npr-19*-expressing neurons was limited to the six non-pharyngeal neuron classes in the head, where AWC is located (ASG, AWA, RIA, RIM, URA, URX). We found that these six classes are presynaptic to 42 different *nape-1,2*-expressing neurons. Approximately half of these neurons receive synaptic input from more than one *npr-19* expressing neuron such that there are 74 coupled pairs fitting the necessary (but not sufficient) anatomical and gene-expression criteria for retrograde AEA signaling. In 14 of these coupled pairs, the postsynaptic neuron is directly presynaptic to AWC, opening the possibility that AWC inherits its AEA sensitivity synaptically.

To test whether classical synaptic pathways render AWC sensitive to AEA, we imaged AWC activity in worms with a null mutation in *unc-13*, the *C. elegans* homolog of Munc13, which is required for exocytosis of the clear-core synaptic vesicles that contain classical neurotransmitters (Richmond et al., 1999). We found that AEA’s effect on food sensitivity in *unc-13* mutants was essentially the same as in wild type worms (Fig. 4B, C; Suppl. Table 4, line 3, 6-7, 9, 13, 15-16, 18). This result makes it unlikely that AWC inherits its AEA sensitivity from synaptic pathways that involve classical neurotransmitters.

In the second indirect model of AEA’s effect on AWC, AEA causes the release of neuromodulators that act on AWC. Most neuromodulatory substances, such as neuropeptides and biogenic amines, are released by exocytosis of dense-core vesicles (Devine & Simpson, 1968; Probert et al., 1983). In mammals, presynaptic terminals that both contain dense-core vesicles and are immunoreactive for the cannabinoid receptor CB1 are a recurring synaptic motif in several brain regions including the CA1 and CA3 of the hippocampus, prefrontal cortex, and basolateral amygdala (Fitzgerald et al., 2019; Takács et al., 2014). To determine whether this motif may be present in *C. elegans*, we used gene expression data (Hammarlund et al., 2018) to search for *npr-19*-expressing neurons that also express *unc-31*, the *C. elegans* ortholog of human CADPS/CAPS, which is required for calcium-regulated dense-core vesicle fusion (Speese et al., 2007). We found that most of the *npr-19*-expressing neurons identified in our study (11 out of 15, Table 1) also express *unc-31*. This result indicates that the anatomical substrate for cannabinoid-mediated release of neuromodulators exists in *C. elegans*.

To test this version of the indirect model, we recorded from AWC in an *unc-31* deletion mutant. If AEA’s effect on AWC were solely the result of neuromodulation mediated by *unc-31*, one would expect this mutation to phenocopy *npr-19* null: exhibiting no AEA effects on AWC responses. This appeared to be the case for the response to favored food, in which there was no effect of AEA (Fig. 4D, E; Suppl. Table 4, line 21, 24-25, 27). AWC responses to non-favored food were still modulated by AEA (Fig. 4D, E; Suppl. Table 4, line 31, 33, 36), but they were increased rather than decreased. The fact that AEA’s modulation of AWC food sensitivity is severely disrupted in *unc-31* mutants supports a model in which NPR-19 receptors activated by AEA promote the release of dense-core vesicles containing modulatory substances that act on AWC.

## Discussion

In mammals, administration of THC or endocannabinoids induces hedonic feeding, meaning an increase in consumption of calorically dense, palatable foods. The present study provides two converging lines of evidence in support of the hypothesis that cannabinoids induce hedonic feeding in *C. elegans*. First, AEA can differentially alter accumulation in favored and non-favored food, causing a larger proportion of worms to accumulate in the former and a smaller proportion in the latter (Fig. 1G). Individual worms tend to exit, explore, and re-enter food patches multiple times over the time scale of our experiments (Shtonda, 2006). Thus, these proportions are mathematically equivalent to the average fraction of time that an individual worm spends feeding on each type of food. Furthermore, worms given an inexhaustible supply of food, feed at a constant rate for at least six hours (Izquierdo et al., 2021), far longer than observation times in this study. Combining these two observations, we can infer that for *C. elegans*, differential accumulation results in differential consumption. Second, AEA differentially alters feeding rate, causing worms to feed at a higher rate in preferred food and a lower rate in non-preferred food (Fig. 1I). Thus, the effect of AEA on feeding rate amplifies its effect on fraction of time feeding in favored and non-favored food patches. The result of this amplification is increased consumption of favored food in a manner consistent with hedonic feeding. We conclude that hedonic feeding is conserved in *C. elegans*.

Our findings confirm and extend previous investigations concerning the role of the endocannabinoid system in regulating feeding in *C. elegans*. The endocannabinoids AEA and 2-AG were previously shown to reduce pumping frequency in animals feeding on nutritionally inferior food (Oakes et al., 2017). We now show that this reduction is part of a broader pattern in which pumping rate on superior food increases and pumping on inferior food decreases. Additionally, we have confirmed that *npr-19* is expressed in a limited number of neurons including the inhibitory pharyngeal motor neuron M3 and the sensory neuron URX. We extend these results by identification of 13 additional *npr-19* expressing neurons including sensory neurons, interneurons, and motor neurons. Of particular interest is the detection of *npr-19* expression in five additional pharyngeal neurons. Thus, 6 of the 20 neurons comprising the pharyngeal nervous system are potential sites for endocannabinoid mediated regulation of pumping rate. It is notable that these six neurons include the motor neuron MC, which is hypothesized to act as the pacemaker neuron for rhythmic pharyngeal contractions (Avery & Horvitzt, 1989; D M Raizen et al., 1995), and M3, which regulates pump duration (Avery, 1993). It will now be important to tackle the question of how pumping rate is modulated in indifferent directions for favored and non-favored foods.

To date, only a small number of studies have examined the effects of cannabinoids on feeding and food preference in invertebrates. Early in evolution, the predominant effect may have been feeding inhibition. Cannabinoid exposure shortens bouts of feeding in Hydra (De Petrocellis et al., 1999). Larvae of the tobacco hornworm moth *Manduca sexta* prefer to eat leaves containing lower rather than higher concentrations of the phytocannabinoid cannabidiol (Park et al., 2019). In adult fruit flies (*Drosophila melanogaster*), pre-exposure to phyto- or endocannabinoids (AEA and 2-AG) for several days before testing reduces consumption of standard food. On the other hand, in side-by-side tests of sugar-yeast solutions with and without added phyto- or endocannabinoids, adult fruit flies prefer the cannabinoid-spiked option. The picture that emerges from these studies is that whereas the original response to cannabinoids may have been feeding suppression, through evolution the opposite effect arose, sometimes in the same organism. As we have shown, *C. elegans* exhibits both increases and decreases in feeding responses under the influence of cannabinoids and does so in a manner that would seem to improve the efficiency of energy homeostasis by promoting consumption of nutritionally superior food and depressing consumption of nutritionally inferior food. At present there is no evidence in mammals for bidirectional modulation of consumption, but our results, together with the logic of homeostasis, predict that such an effect may exist under certain conditions.

Although administration of cannabinoids causes hedonic feeding in *C. elegans* and mammals, there are notable differences in how it is expressed. One experimental design commonly used in mammalian studies is to measure consumption of a single test food, which is either standard lab food or a more palatable food. In such experiments, consumption of both food types is increased (Williams et al., 1998; Williams & Kirkham, 1999). The analogous experiment in the present study is the experiment of Fig. 1I, in which consumption (inferred from pumping rate) was measured in response to either favored or non-favored food. We found that consumption of favored food increases as in mammalian studies whereas, in contrast, consumption of non-favored food decreases. A second experimental design commonly used in mammalian studies is to measure consumption of standard and palatable foods when the two foods are presented together. In this type of experiment, cannabinoids increase consumption of palatable food, but consumption of standard food is unchanged (Brown et al., 1977; Deshmukh & Sharma, 2012; Escartín-Pérez et al., 2009b; Koch & Matthews, 2001; Shinohara et al., 2009). Cannabinoid receptor antagonists produce the complementary effect: reduced consumption of palatable food with little or no change in consumption of standard food. The analogous experiments in the present study are the T-maze assays in which maze arms are baited with favored and non-favored food. We find that following cannabinoid administration, consumption of favor food increases whereas consumption of non-favored food decreases.

Thus, considering both experimental designs, the effects of cannabinoid exposure on consumption in *C. elegans* are bidirectional, whereas in mammals they are not. It is conceivable that a bidirectional response is advantageous in that it produces a stronger bias in favor of superior food than a unidirectional response, raising the question of why bidirectional responses have not been reported in mammals. There are, of course, considerable differences in the feeding ecology of nematodes and mammals; perhaps mammals evolved under a different set of constraints under which unidirectional responses are the better strategy. On the other hand, differences in experimental procedures may explain the absence of bidirectional responses. For example, in mammalian studies in which the two foods are presented together, standard and palatable foods are placed in close proximity within a small cage, with the result that there is essentially no cost in terms of physical effort for the animal to switch from one feeding location to the other. It is conceivable that increasing the switching cost (Salamone et al., 1994) could lead to a differential effect on consumption in mammals.

We propose the following model of differential accumulation on food leading to hedonic feeding in *C. elegans*. The model focusses on the olfactory neuron AWC, which is necessary and sufficient for navigation to the source of food-related odors (Kocabas et al., 2012) and exhibits bidirectional modulation by AEA. Calcium imaging shows that AWC is activated by food removal, regardless of whether favored or non-favored food is removed (Fig. 2C)(Chalasani et al., 2007). Previous studies have demonstrated that exogenous activation of AWC triggers two previously described behavioral motifs known to contribute to locomotion oriented toward attractive odors. First, its activation truncates bends of the head and neck that occur during the worm’s normal sinusoidal locomotion (Kocabas et al., 2012). This means that each time a body bend moves the head away from an odor source, AWC will activate, and this bend will be truncated. Over time, successive truncations of bends in the wrong direction steer the animal in the right direction: toward the odor source; this widely conserved behavioral motif is known as *klinotaxis* (Fraenkel & Gunn, 1961). Second, activation of AWC causes the animal to stop moving forward, reverse, and resume locomotion in a new direction that is better aligned with the food odor source (Gordus et al., 2015; Gray et al., 2005); this behavioral motif is known as a *pirouette* (Pierce-Shimomura et al., 1999). Both motifs not only promote navigation toward a patch of food, but also promote retention in a patch. For example, a pirouette initiated when the worm’s head protrudes beyond the food-patch boundary will return the worm into the patch. We find that AEA exposure increases AWC’s response to the removal of favored food (Fig. 2C). In the proposed model, this effect both accentuates klinotaxis and increases the probability of pirouettes caused by locomotion away from the odor source. The net result is enhanced approach to, and retention in, patches of favored food. Conversely, we also find that AEA exposure decreases responses to removal of non-favored food. This effect weakens klinotaxis and decreases pirouette probability, resulting in diminished approach and retention in non-favored food. The result of these two processes is increased or decreased accumulation, respectively, in patches of favored and non-favored food.

The requirement for *ceh-36* in rendering *C. elegans* food preferences sensitive to AEA (Fig. 2B) suggests that AWC neurons provide a necessary link between AEA and hedonic feeding. However, this experiment does not have statistical power sufficient to rule out contributions from other chemosensory neurons. Of particular interest are two chemosensory neurons AWA and ASG, both of which express *npr-19* (Table 1) and are required for chemotaxis (Bargmann et al., 1993; Bargmann & Horvitz, 1991). It will now be important to map cannabinoid sensitivity across the entire population or food-sensitive odors to understand how cannabinoids alter the overall chemosensory representation of favored and non-favored foods.

Cannabinoids have been observed to modify chemosensitivity at several levels in mammals. Both AEA and 2-AG amplify the response of primary chemosensory cells, such as the sweet-taste cells in the tongue (Yoshida et al., 2010, 2013), which may help to explain increased consumption of sweet foods and liquids. Cannabinoids can also increase the sensitivity of the mammalian olfactory system as measured during food-odor exploration (Heinbockel & Straiker, 2021; Nogi et al., 2020; Soria-Gómez et al., 2014). We observed an analogous effect in *C. elegans*, in that AEA alters the sensitivity of a primary chemosensory neuron, AWC. In unexposed worms, AWC is equally sensitive to favored and non-favored food, suggesting it cannot detect a difference in the odors released by the two food types. However, in remarkable alignment with the observed bidirectional changes in food preference in worms exposed to AEA, this neuron becomes more sensitive to favored food and less sensitive to non-favored food, therefore acquiring the ability to discriminate between the odors of these foods.

AEA’s effect on AWC appears to be indirect. Our results are consistent with a model in which AEA activates NPR-19 receptors to promote release of dense-core vesicles containing neuromodulators that act on AWC. This model is supported by evidence in *C. elegans* that 2-AG, which is capable of activating NPR-19, stimulates widespread release of serotonin (Oakes et al., 2017, 2019); thus, NPR-19 activation seems capable of promoting dense-core vesicle release. Additionally, AWC expresses receptors for biogenic amines, and it responds to neuropeptides released by neighboring neurons (Chalasani et al., 2010; Leinwand & Chalasani, 2013), suggesting that it has postsynaptic mechanisms for responding to neuromodulation. Identification of one or more neuromodulators responsible for AEA’s effect on AWC, together with their associated receptors, will be an important step in answering the question of how AEA causes differential changes in food-odor sensitivity.

Our results establish a new role for endocannabinoids in *C. elegans*: the induction of hedonic feeding. There is general agreement that the endocannabinoid system and its molecular constituents offer significant prospects for pharmacological management of health, including eating disorders and substance abuse (Parsons & Hurd, 2015). Clear parallels between the behavioral, neuronal, and genetic basis of hedonic feeding in *C. elegans* and mammals establish the utility of this organism as a new genetic model for the investigation of molecular and cellular basis of these and related disorders.

## Materials and Methods

### Strains

Animals were cultivated under standard conditions (Brenner, 1974) using *E. coli* OP50 as a food source. Young adults of the following strains were used in all experiments:

| Experiment | Strain | Genotype |
| --- | --- | --- |
| Reference strain | N2, Bristol | Wild type |
| Preference and feeding assays | FK311 | *ceh-36*(ks86) |
|  | RB1668 | *npr-19*(ok2068) |
|  | XL324 | ntIS1701[*npr-19*::CNR1::gfp-*npr-19*(1.1);*unc-122*::RFP] |
|  | XL325 | ntIS1702[*npr-19*::*npr-19*::gfp-*npr-19*(1.1)] |
| Calcium imaging | XL322 | ntIS1703[*str-2*::GCaMP6::wCherry;*unc-122*::dsRed] |
|  | XL327 | *unc-13*(e51);ntIS1703[*str-2*::GCaMP6::wCherry;*unc-122*::dsRed] |
|  | XL326 | *unc-31*(e928);ntIS1703[*str-2*::GCaMP6::wCherry;*unc-122*::dsRed] |
|  | XL346 | *npr-19*(ok2068);ntIS1912[*str-2*::GCaMP6::wCherry;*unc-122*::dsRed] |
| *npr-19* expression pattern | XL334 | otIS544[*cho-1*::SL2::mCherry::H2B+*pha-1*(+);ntIS19114[*npr-19*::GFP1.1;*unc-122*::dsred] |
|  | XL335 | ntIS19114[*npr-19*::GFP1.1;*unc-122*::dsred];otIS518[*eat-4*::SL2::mCherry::H2B+*pha-1*(+)] |

### Bacteria

The following streptomycin-resistant bacterial strains were used in this study: DA1885 (*Bacillus simplex*), DA1877 (*Comamonas* sp.), *E. Coli* HB101, and *E. Coli* DA837. Bacteria were grown overnight at 37°C in presence of 50 mg/ml streptomycin, concentrated by centrifugation, rinsed three times with either M9 medium (for EPG experiments) or A0 buffer (for behavioral/imaging experiments; MgSO_4_ 1 mM, CaCl_2_ 1 mM, HEPES 10 mM, glycerol to 350 mOsm, pH 7), and resuspended to their final concentration. Concentration was defined as optical density at 600 nm (OD_600_), as measured with a DSM cell density meter (Laxco, Bothell, WA, USA). All measurements were performed on samples diluted into the linear range of the instrument (OD 0.1-1). Previous experiments determined that OD_600_ = 1 corresponds to approximately 2.35 × 10^9^ and 2.00 × 10^9^ colony forming units/mL of *Comamonas* and *Simplex*, respectively (Katzen *et al*., 2021).

### Animal preparation

Worms were washed five times in M9 for EPG experiments or A0 buffer (see above) for behavioral/imaging experiments. Worms were then incubated for 20 minutes with either background solution alone or background solution + 300 μM (electropharyngeogram experiments) or 100 μM (behavioral assays and calcium imaging experiments) Arachidonoylethanolamide (AEA, Cayman chemical, Ann Arbor, MI, USA). The incubation time and relatively high concentration reflects the low permeability of the *C. elegans* cuticle to exogenous molecules (Rand & Johnson, 1995; Sandhu et al., 2021).

### Behavioral assays

Freshly poured NGM agar plates were dried in a dehydrator for 45 minutes at 45°C. A maze cut from foam sheets (Darice, Strongsville, OH, USA) using a laser cutter was placed on each plate (Fig. 1A). Maze arms were seeded with 4.5 μl of bacteria. Animals were deposited at the starting point of the maze by liquid transfer and a transparent plastic disc was placed over the maze to eliminate air currents; 12 plates were placed on a flatbed scanner and simultaneously imaged every 15 minutes (Mathew et al., 2012; Stroustrup et al., 2013). The number of worms in the two patches of food and the region between them was counted manually and a preference index *I* calculated as: *I* = (*n*_F_ − *n*_NF_)/(*n*_F_ + *n*_NF_), where *n*_F_ is the number of worms in the favored food patch, and *n*_NF_ is the number of worms in the non-favored food patch. Worms that did not leave the starting point were excluded. For experiments involving mutants, a cohort of N2 animals was run in parallel on the same day. Data from statistically indistinguishable N2 cohorts were pooled where possible. In some experiments, a paralytic agent (sodium azide, NaN_3_, 3 μl at 20 mM), was added to each food patch to prevent animals from leaving the patch of food after reaching it. Sodium azide diffuses through the agar over time and its action is not instantaneous. These two characteristics resulted in some worms becoming paralyzed around rather than in the patch of food, as they stop short of the patch or escape the patch briefly before becoming paralyzed. To account for these effects all worms within 5mm of the end of the maze’s arm, rather than on food, were used when calculating preference index.

### Electropharyngeograms

Pharyngeal pumping was measured electrophysiologically (Lockery et al., 2012) using a ScreenChip microfluidic system (InVivo Biosystems, Eugene, OR, USA). Briefly, following pre-incubation as described above, worms were loaded into the worm reservoir of the microfluidic device which was pre-filled with bacterial food (OD_600_ = 0.8) ±AEA 300 μM; this food density was chosen to reduce possible ceiling effects on pumping rate modulation by AEA. To record voltage transients associated with pharyngeal pumping (David M. Raizen & Avery, 1994)., worms were transferred on at a time from the reservoir to the recording channel of the device such that the worm was positioned between a pair of electrodes connected to a differential amplifier. Worms were given three minutes to acclimate to the channel before and recorded for one minute. Mean pumping frequency was extracted using custom code written in Igor Pro (Wavemetrics, Lake Oswego, OR, USA).

### Calcium imaging

After pre-incubation with buffer or buffer +AEA (see: animal preparation), worms were immobilized in a custom microfluidic chip and presented with alternating 30-second epochs of buffer and bacteria (either *B. Simplex* or *Comamonas sp*. at OD_600_ 1, at a flow rate of 100 μl/min) for 3 minutes. Optical recordings of GCaMP6-expressing AWC neurons were performed on a Zeiss Axiovert 135, using a Zeiss Plan-Apochromat 40× oil, 1.4 NA objective, a X-Cite 120Q illuminator, a 470/40 excitation filter, and a 560/40 emission filter. Neurons were imaged at 3-10 Hz on an ORCA-ERA camera (Hamamatsu). Images were analyzed using custom code written in MATLAB: the change in fluorescence in a hand-drawn region of interest that contained only the soma and neurite. Data were normalized to the average fluorescence *F*_o_ computed over the 15 second interval before the first food stimulus. We computed normalized fluorescence change as Δ*F*(*t*)/*F*_o_, where Δ*F*(*t*) = *F*(*t*) − *F*_o_; following convention, we refer to this measure as “Δ*F*/*F*.” For comparison of treatment groups, we used the peak amplitude of post-stimulus Δ*F*/*F*. In some animals, AWC appeared not to respond to the food stimulus, regardless of treatment group. To classify particular AWC neurons as responsive or non-responsive, we obtained the distribution of peak Δ*F*/*F* values in control experiments in which the stimulus channel contained no food; responsive neurons were defined as those whose peak Δ*F*/*F* value exceeded the 90^th^ percentile of this distribution. Critically, the percentage of non-responders did not vary between AEA-treated animals and (25.46% vs 22.49% respectively; *χ*^2^(1,759) = 0.699, *p* = 0.4031).

### Expression profile for *npr-19*

Worms were immobilized with 10 mM sodium azide (NaN_3_) and mounted on 5% agarose pads formed on glass slides. Image stacks (30-80 images) were acquired using a Zeiss confocal microscope (LSM800, ZEN software) at 40X magnification. Identification of neurons was done based on published expression profiles of the p*cho-1*::mCherry (Pereira et al., 2015) and p*eat-4*::mCherry (Serrano-Saiz et al., 2013) transgenes in *C. elegans*. Individual neurons were identified by mCherry expression and the relative positions of their cell bodies; *npr-19* expression was visualized using a p*npr-19*::GFP transgene. Co-expression of GFP and mCherry was assessed by visual inspection using 3D image analysis software Imaris (Oxford Instruments). Representative images (Fig. 4A) are maximum intensity projections of 30-80 frames computed using ImageJ software (Collins, 2007). Expression of the NPR-19 receptor was widespread in body wall muscles, but restricted to 29 neurons in the head (27 - 31, 95% confidence interval, *n* = 20 worms imaged) and 8 neurons in the tail (7.8 - 8.5, 95% confidence interval, *n* = 22 worms imaged) (Suppl. Table 6). Overall, 28 of the *npr-19*-expressing neurons co-localized with either *cho-1* or *eat-4*, whereas ∼9 did not co-localize with either marker. The identity of the latter cells was ascertained based on cell body position and morphology, and verified by *npr-19* expression (threshold = 2) as reported in the *C. elegans* Neuronal Gene Expression Map & Network (CeNGEN) consortium database (Hammarlund et al., 2018).

### Statistics

A detailed description of statistical tests used, their results, and their interpretation is presented in Supplemental Tables 1-5. Data were checked for normality with a Kolmogorov-Smirnov test.

#### Number of replicates

The minimal sample size for the T-maze assays were based on pilot experiments which showed an acceptable effect size with ∼10 replicates per experimental condition. Similarly, the minimal number of replicates for EPG experiments and imaging experiments were based on previously published data in which mutants/treatments could be distinguished with ∼10 replicates.

#### Effect sizes

Effect sizes were computed as follow: Cohen’s *d* for *t*-tests, partial eta-squared for ANOVAs, and |*z*|/√*n* for Mann-Whitney test, where *z* is the *z*-score and *n* is the number of observations.

#### Behavioral experiments (T-mazes)

Preference indices were analyzed using a two-factor ANOVA with repeated measures (effect of AEA × effect of time, with time as a repeated measure). For easier presentation, an average index across the four time-points was calculated and displayed (Fig. 1C-F, 2B, 3A). All time series are nonetheless available for inspection in Fig. 1A, 2A and Supplemental Fig. 1 and 2. The effect of AEA was deemed significant if main effect of AEA was significant in the ANOVA. Averaging the four time points in a series would only be problematic if there was a non-ordinal interaction AEA × time. Inspection of ANOVA results and time series reveal that the only AEA × time interaction in Fig. 1E is ordinal and minimal. In cases where the effect of time was important (Fig. 2A) or the interaction AEA × time was meaningful (Fig. 1G) the time series of preference indices was presented. The comparison of preference indices between N2 and mutants relied on a two-factor ANOVA (effect of strain × effect of AEA). The average preference index across the four time-points was used for the comparison. In addition to an ANOVA, planned comparisons were incorporated in the experimental design using t-tests and focusing on four scientifically relevant contrasts: (1) mutants, AEA– vs AEA+; (2) N2, AEA– vs AEA+; (3) AEA–, mutants vs N2; (4) AEA+, mutants vs N2.

#### Electropharyngeograms

As the data were not normally distributed in most of the cohorts, a non-parametric test (Mann-Whitney) was used to compared pumping frequencies between strains/treatments.

#### Calcium imaging

Peak ΔF/F was used as the primary measure. A two-factor ANOVA (effect of AEA × effect of bacteria type) was used to assess the effect of AEA on AWC responses. Planned t-tests were focused on four contrasts: (1) favored food, AEA– vs AEA+; (2) non-favored food, AEA– vs AEA+; (3) AEA-, favored food vs non-favored food; (4) AEA+, favored food vs non-favored food. For comparisons between N2 and mutants, a two-factor ANOVAs (effect of AEA × effect of strain) was performed for each of the bacteria type (favored and non-favored) and followed by four contrasts (t-tests): (1) mutants, AEA– vs AEA+; (2) N2, AEA– vs AEA+; (3) AEA–, mutants vs N2; (4) AEA+, mutants vs N2.

#### Multiple comparisons

No correction for multiple comparisons was applied in *t*-tests used in pair-wise comparisons of means in multifactor experiments as the experimental design in this study relied on a small number (3 per condition) of planned (a priori), rather than unplanned (a posteriori), scientifically relevant contrasts (Keppel & Zedeck, 1989).

## Acknowledgements

We thank Richard Komuniecki for the *npr-19-*null and rescue worm strains. The *unc-13, unc-31, ceh-36, cho-1*, and *eat-4* worm strains were provided by the CGC, which is funded by NIH Office of Research Infrastructure Programs (P40 OD010440). We also would like to thank Oliver Hobert, Jonathan Millet, and Jon Pierce for thoughtful discussion of the project. We thank Chris Doe for use of his Zeiss LSM800 confocal microscope for imaging. Finally, we would like to thank Kathy Chicas-Cruz for constructing our calcium imaging strains. Funding for this project was provided by NIDA (DA047645) and NIGM (GM129576).

## Supplemental material

**Supp. Fig 1.**
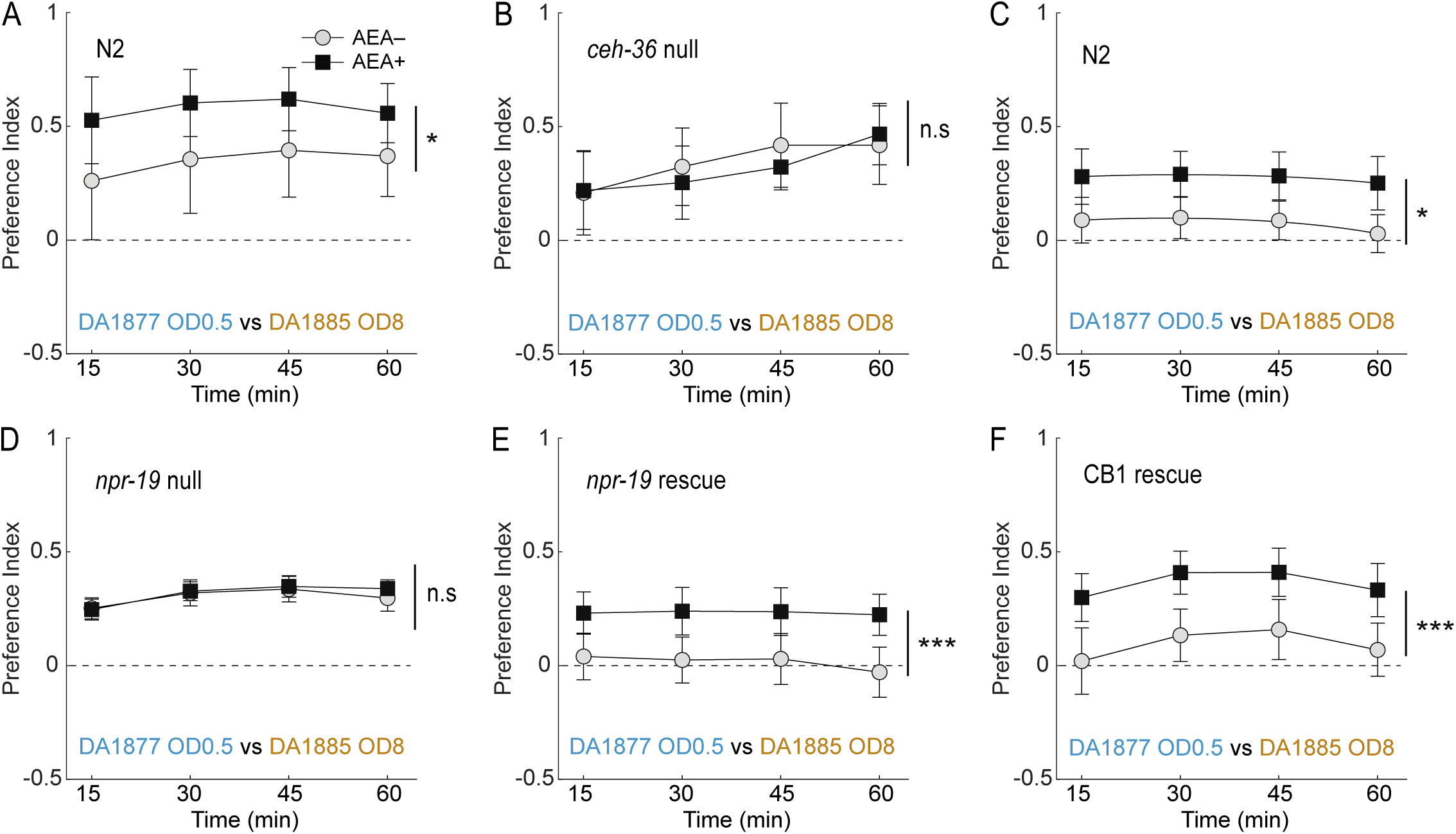
Effect of baseline preference and bacteria identity on preference time course. Mean preference index (*I*) versus time for AEA-exposed animals (AEA+) and unexposed controls (AEA–), where *I* > 0 is preference for favored food, *I* < 0 is preference for non-favored food, and *I* = 0 is indifference (*dashed line*). **A**. Time course, Fig. 1D. **B**. Time course, Fig. 1E. **C**. Time course, Fig. 1F. For statistics in **A**-**C**, see Supp. Table 1. Symbols: *, *p* < 0.05; **, *p* < 0.01; n.s., not significant. Error bars, 95% confidence intervals.

**Supp. Fig 2.**
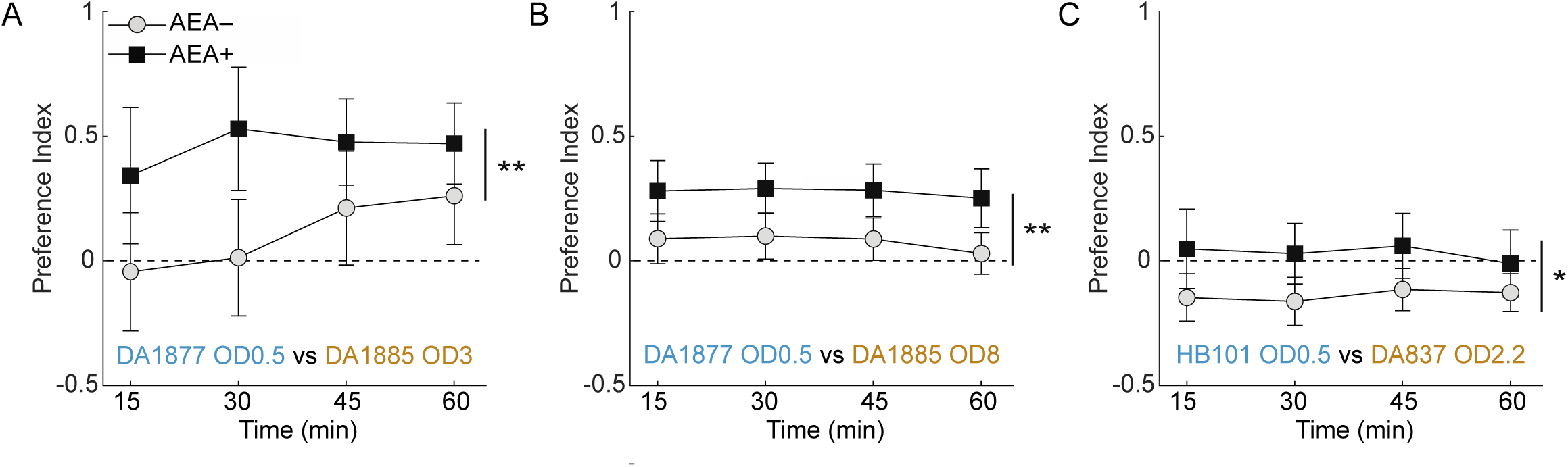
Effect of genetic background on preference time course. Mean preference index (*I*) versus time for AEA-exposed animals (AEA+) and unexposed controls (AEA–), where *I* > 0 is preference for favored food, *I* < 0 is preference for non-favored food, and *I* = 0 is indifference (*dashed line*). **A**. Time course, Fig. 2B, N2. **B**. Time course, Fig. 2B, *ceh-36*. **C**. Time course, Fig. 3A, N2. **D**. Time course, Fig. 3A, *npr-19* null. **E**. Time course, Fig. 3A, *npr-19* rescue. **F**. Time course, Fig. 3A, CB1 rescue. **A**-**F**. For statistics, see Supp. Tables 2, 3. Symbols: *, *p* < 0.05; ***, *p* < 0.001; n.s., not significant. Error bars, 95% confidence intervals.

**Supp. Fig 3.**
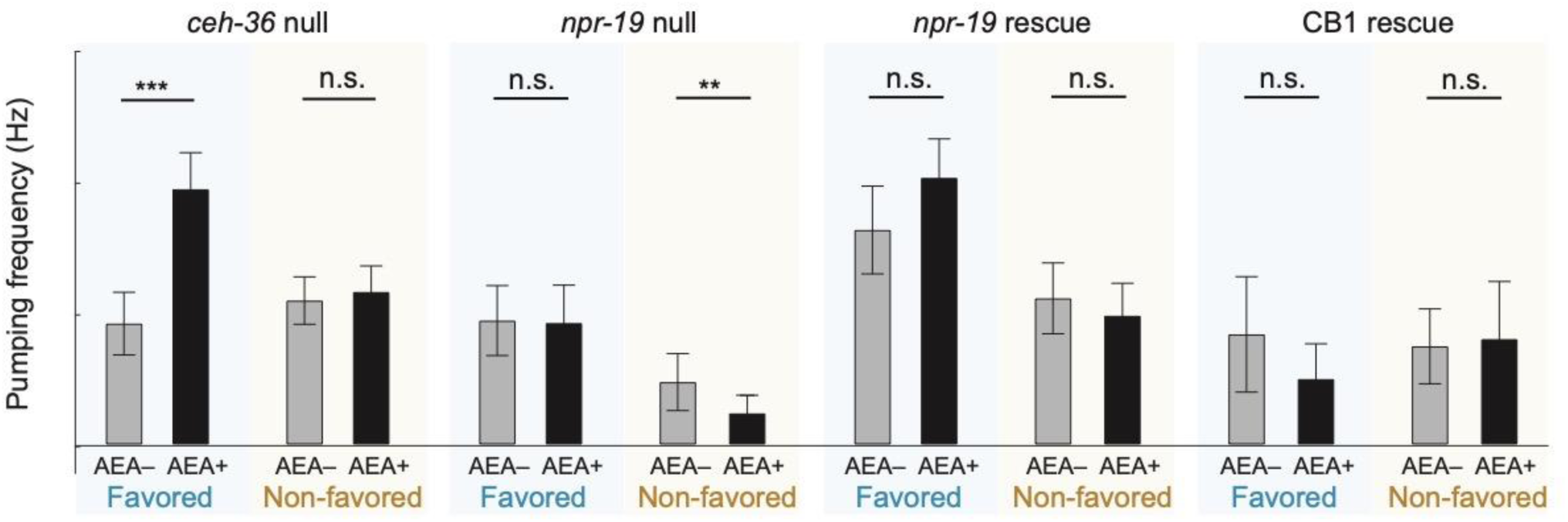
Effect of AEA on pharyngeal pumping frequency in different genetic backgrounds. Mean pumping frequency in favored and non-favored food is shown for or AEA-exposed animals (AEA+) and unexposed controls (AEA–). Favored food, DA1877, OD 0.8; non-favored food, DA1885, OD 0.8. For statistics, see Supp. Table 5. Symbols: **, *p* < 0.01; ***, *p* < 0.001. Error bars, 95% confidence intervals.

**Supp. Fig 4.**
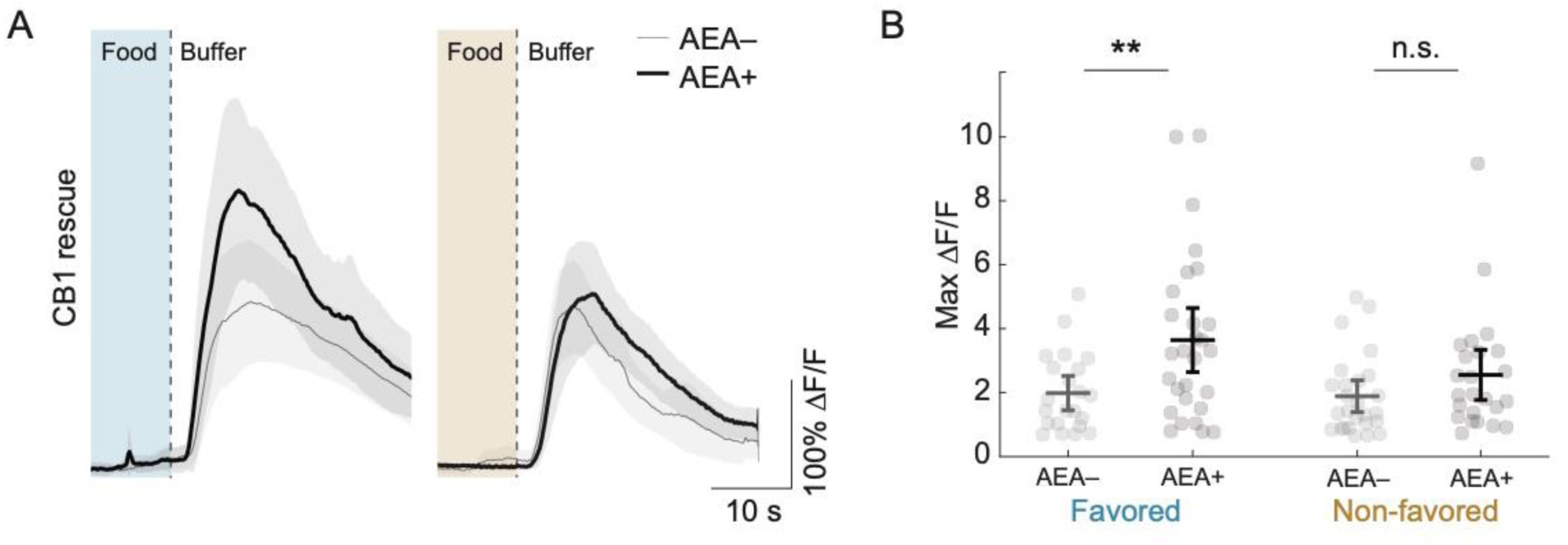
CB1 partial rescue of AEA sensitivity in AWC neurons. **A**. Effect of AEA on the response of AWC neurons to the removal of favored or non-favored food in *npr-19* mutants in which CB1 was overexpressed under control of the *npr-19* promoter. Each trace is average normalized fluorescence change (Δ*F*/*F*) versus time. Favored food (blue), DA1877, OD 1; non-favored food (orange), DA1885, OD 1. **B**. Summary of the data in **A**, showing mean peak Δ*F*/*F*. For statistics in **A**-**B**, see Supp. Table 5. Symbols: **, *p* < 0.01. Error bars or shading, 95% confidence intervals.

**Supplemental Table 1.**
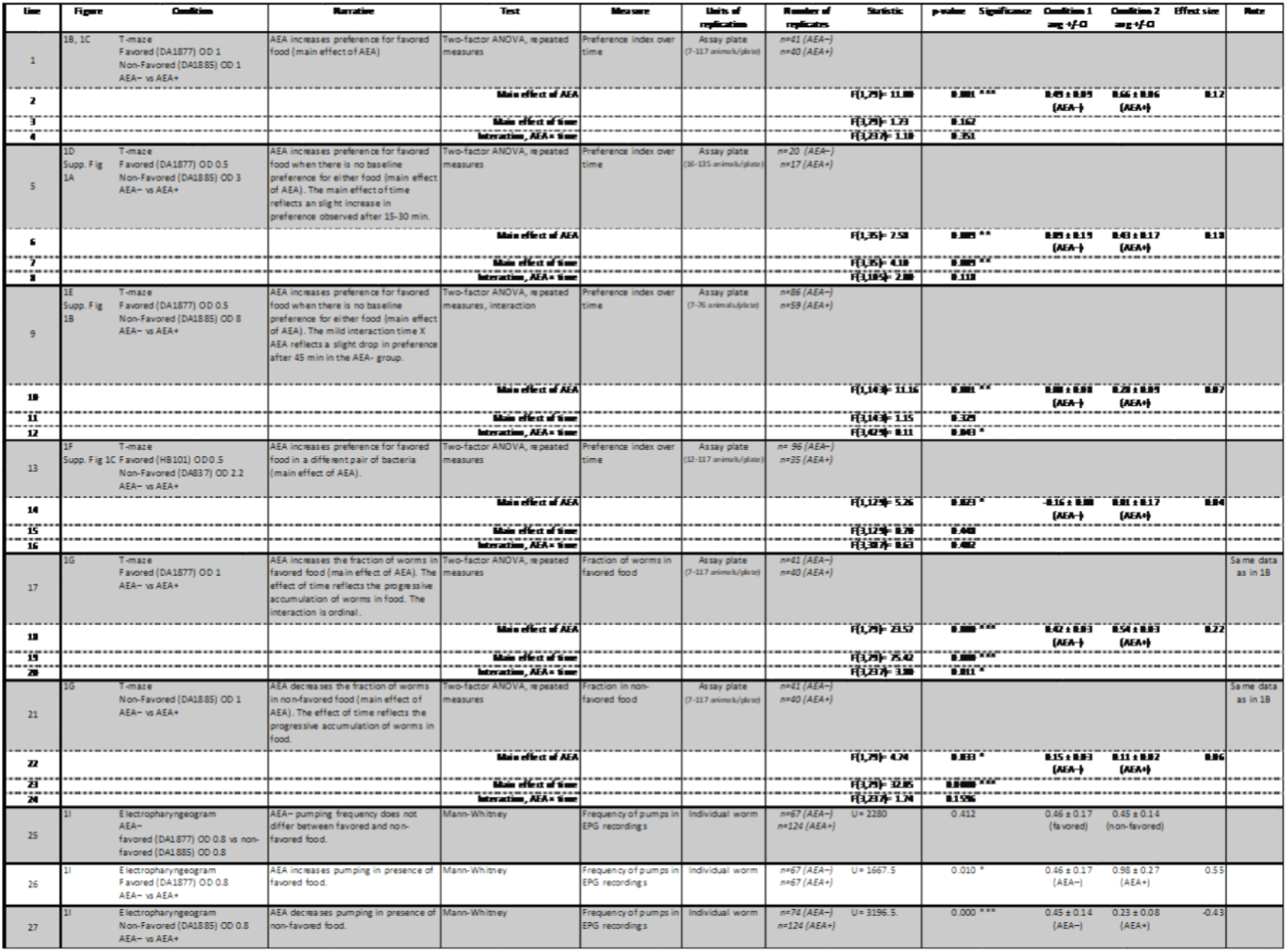
Statistics for Fig. 1 and Supp. Fig. 1. Experimental conditions and comparisons tested are described in column 3. Stars in the Significance column indicate significance levels: *, *p* < 0.05; **, *p* < 0.01; ***, *p* < 0.001. Effect sizes were computed as described in Materials and Methods and 95% confidence intervals were used as a dispersion measure.

**Supplemental Table 2.**
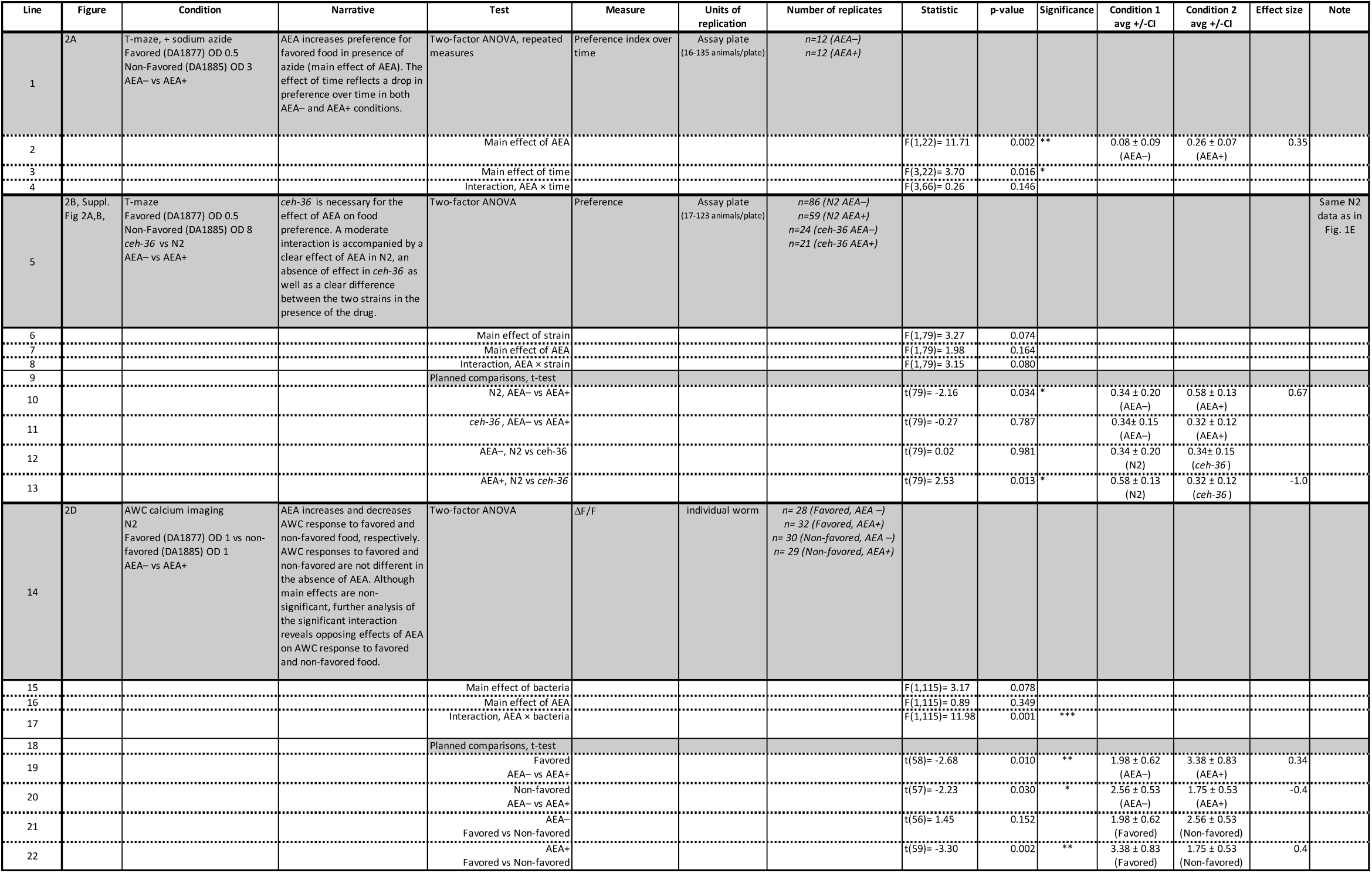
Statistics for Fig. 2 and Supp. Fig. 2 A, B. Experimental conditions and comparisons tested are described in column 3. Stars in the Significance column indicate significance levels: *, *p* < 0.05; **, *p* < 0.01; ***, *p* < 0.001. Effect sizes were computed as described in Materials and Methods and 95% confidence intervals were used as a dispersion measure.

**Supplemental Table 3.**
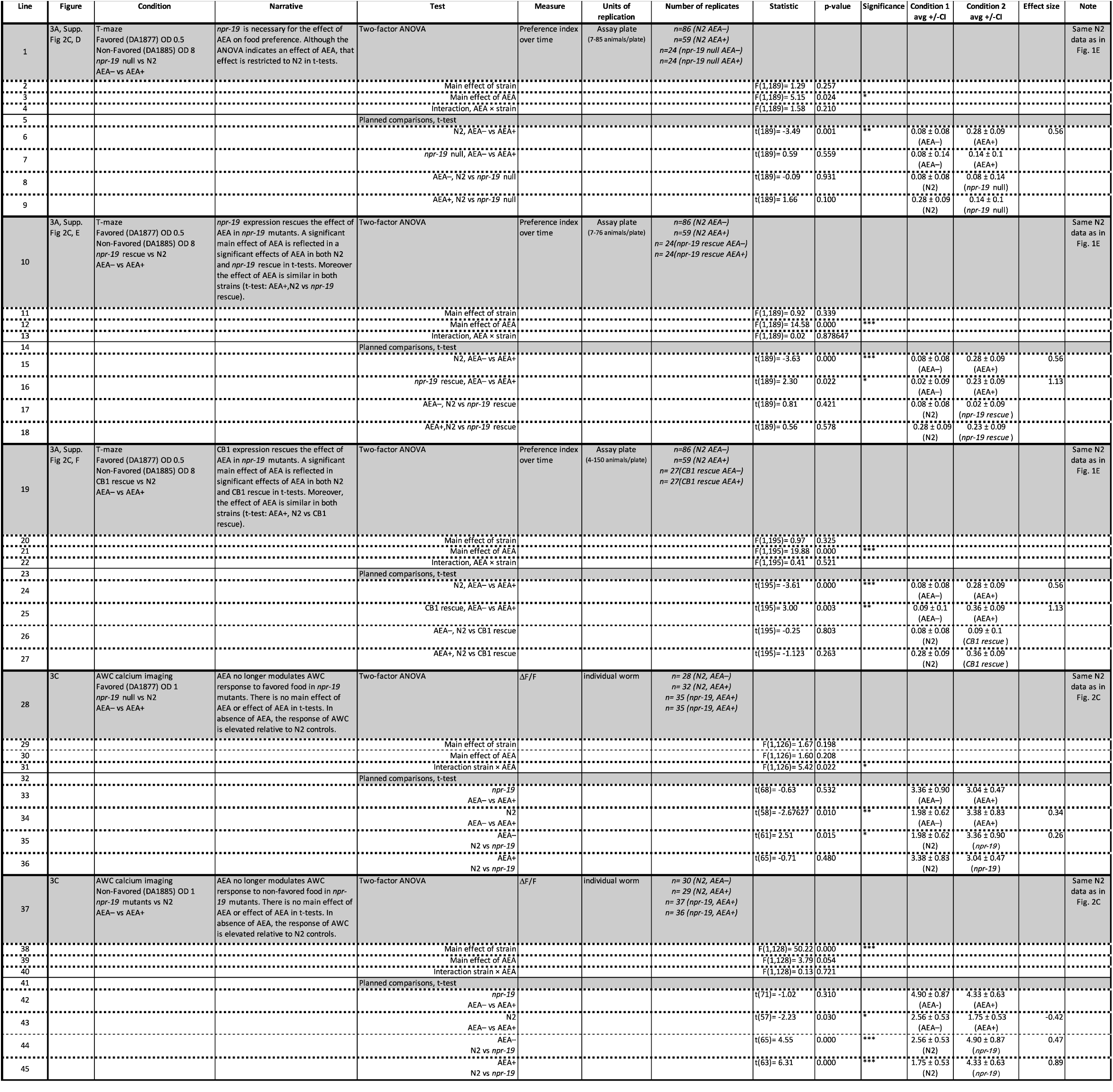
Statistics for Fig. 3 and Supp. Fig. 2 C-F. Experimental conditions and comparisons tested are described in column 3. Stars in the Significance column indicate significance levels: *, *p* < 0.05; **, *p* < 0.01; ***, *p* < 0.001. Effect sizes were computed as described in Materials and Methods and 95% confidence intervals were used as a dispersion measure.

**Supplemental Table 4.**
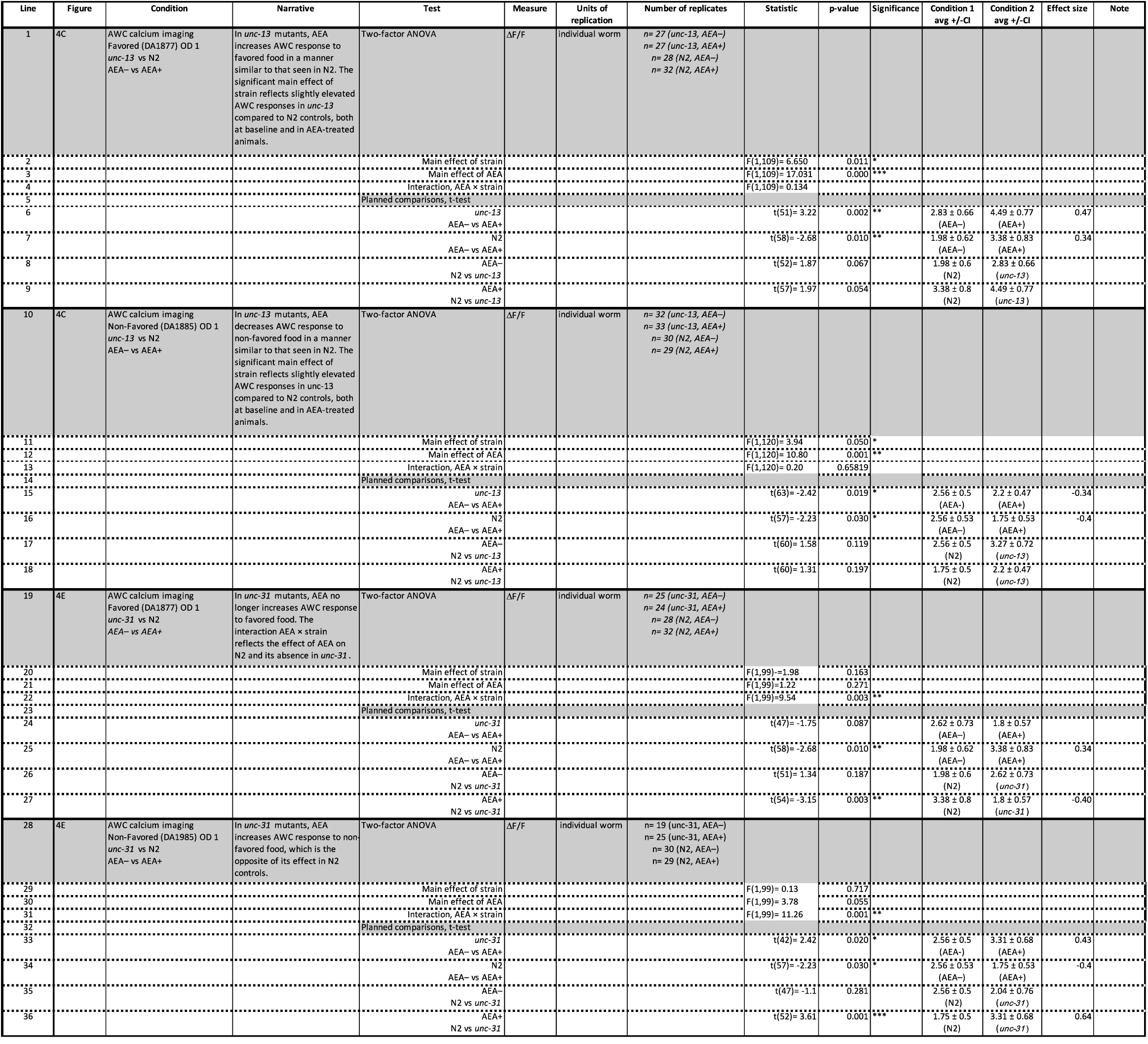
Statistics for Fig. 4. Experimental conditions and comparisons tested are described in column 3. Stars in the Significance column indicate significance levels: *, *p* < 0.05; **, *p* < 0.01; ***, *p* < 0.001. Effect sizes were computed as described in Materials and Methods and 95% confidence intervals were used as a dispersion measure.

**Supplemental Table 5.**
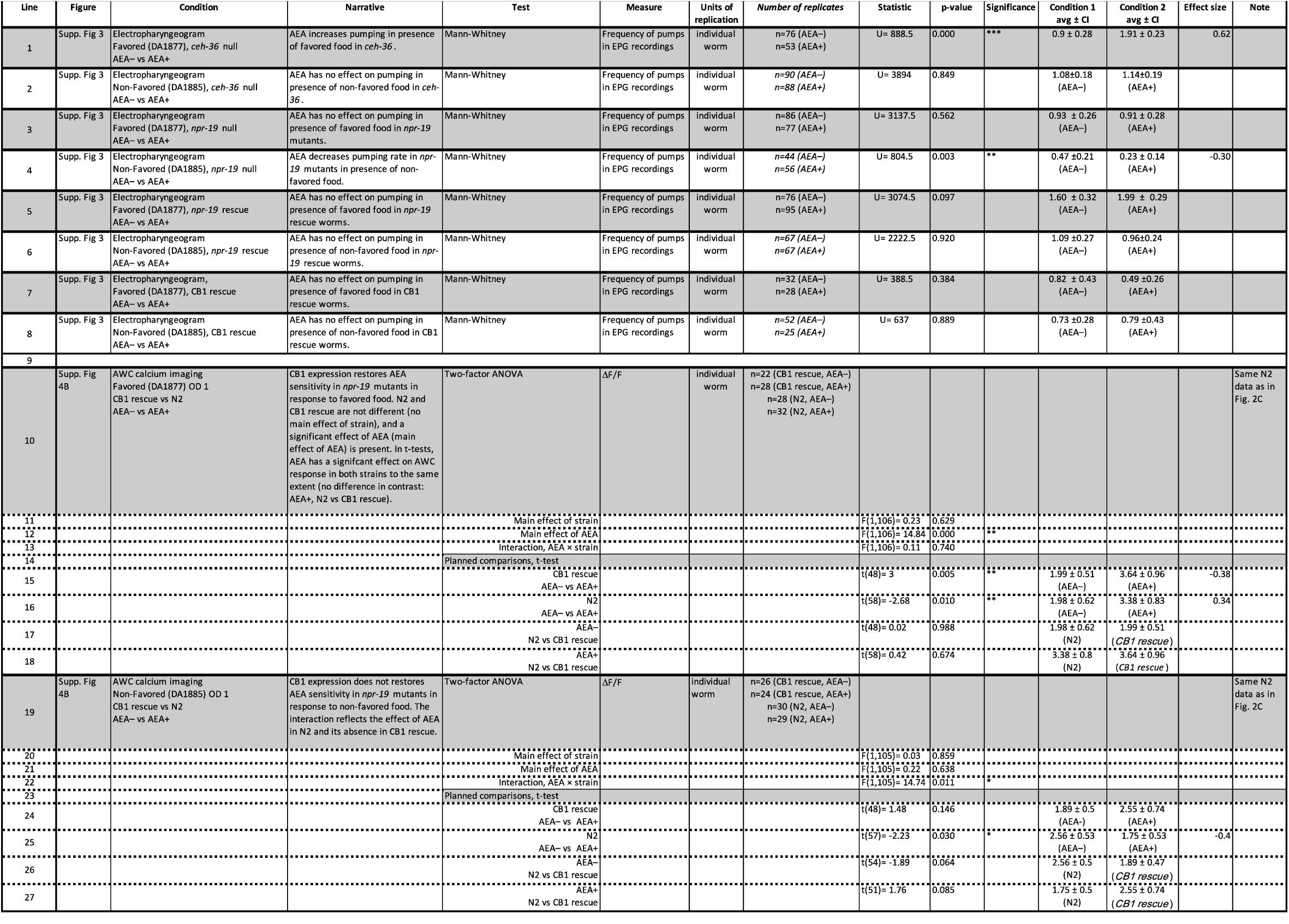
Statistics for Supp. Fig. 3, Supp. Fig. 4. Experimental conditions and comparisons tested are described in column 3. Stars in the Significance column indicate significance levels: *, *p* < 0.05; **, *p* < 0.01; ***, *p* < 0.001. Effect sizes were computed as described in Materials and Methods and 95% confidence intervals were used as a dispersion measure.

**Supplemental Table 6.**
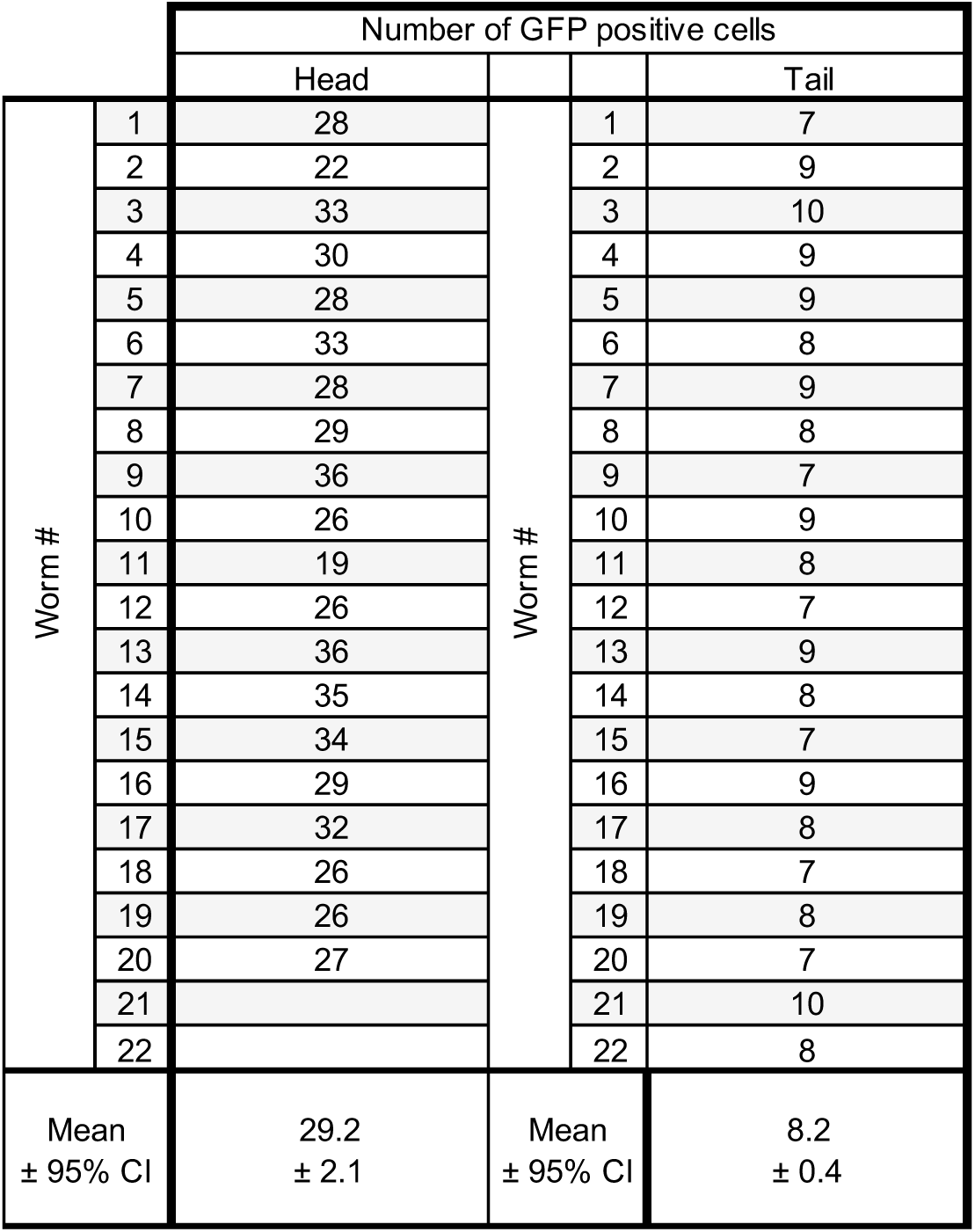
Counts of *npr-19*-expressing neurons in the head and tail. Number of p*npr-19*::GFP positive neurons present in the head (*n* = 20 worms), or the tail (*n* = 22 worms).

|  |  | Number of GFP positive cells |  |  |  |
| --- | --- | --- | --- | --- | --- |
|  |  | Head |  |  | Tail |
| Worm # | 1 | 28 | Worm # | 1 | 7 |
|  | 2 | 22 |  | 2 | 9 |
|  | 3 | 33 |  | 3 | 10 |
|  | 4 | 30 |  | 4 | 9 |
|  | 5 | 28 |  | 5 | 9 |
|  | 6 | 33 |  | 6 | 8 |
|  | 7 | 28 |  | 7 | 9 |
|  | 8 | 29 |  | 8 | 8 |
|  | 9 | 36 |  | 9 | 7 |
|  | 10 | 26 |  | 10 | 9 |
|  | 11 | 19 |  | 11 | 8 |
|  | 12 | 26 |  | 12 | 7 |
|  | 13 | 36 |  | 13 | 9 |
|  | 14 | 35 |  | 14 | 8 |
|  | 15 | 34 |  | 15 | 7 |
|  | 16 | 29 |  | 16 | 9 |
|  | 17 | 32 |  | 17 | 8 |
|  | 18 | 26 |  | 18 | 7 |
|  | 19 | 26 |  | 19 | 8 |
|  | 20 | 27 |  | 20 | 7 |
|  | 21 |  |  | 21 | 10 |
|  | 22 |  |  | 22 | 8 |
| Mean<br>± 95% CI |  | 29.2<br>± 2.1 | Mean<br>± 95% CI |  | 8.2<br>± 0.4 |

